# Elucidating Collective Translocation of Nanoparticles Across the Skin Lipid Barrier: A Molecular Dynamics Study

**DOI:** 10.1101/2022.01.20.477051

**Authors:** Yogesh Badhe, Pradyumn Sharma, Rakesh Gupta, Beena Rai

**Author notes:** contributed equally (Yogesh Badhe, Pradyumn Sharma, Rakesh Gupta).

## Abstract

The top layer of skin, the stratum corneum, provides a formidable barrier to the skin. Nanoparticles are utilized and further explored for personal and health care applications related to the skin. In past years several researchers have studied the translocation and permeation of nanoparticles of various shapes, sizes, and surface chemistry through the cell membranes. Most of these studies focused on a single nanoparticle and a simple bilayer system, whereas skin has a highly complex lipid membrane architecture. Moreover, it is highly unlikely that a nanoparticle formulation applied on the skin will not have multiple nanoparticle-nanoparticle and skin-nanoparticle interactions. In this study, we have utilized coarse-grained MARTINI molecular dynamics simulations to assess the interactions of two types (bare and dodecane-thiol coated) of nanoparticles with two models (single bilayer and double bilayer) of skin lipid membranes. The nanoparticles were found to be partitioned from the water layer to the lipid membrane as an individual entity as well as in the cluster form. It was discovered that each nanoparticle reached the interior of both single bilayer and double bilayer membrane irrespective of nanoparticle type and concentration, though coated particles were observed to efficiently traverse across bilayer when compared with bare particles. The coated nanoparticles also created a single large cluster inside the membrane, whereas bare nanoparticles were found in small clusters. Both the nanoparticles exhibited preferential interactions with cholesterol molecules present in the lipid membrane as compared to other lipid components of the membrane. We have also observed that the single membrane model exhibited unrealistic instability at moderate to the higher concentration of nanoparticles, and hence for translocation study, at minimum double bilayer model should be employed.

## 1 Introduction

The outermost layer of the skin, stratum corneum (SC), provides barrier against chemical, biological and mechanical stimuli [1,2]. This layer allows only small and relatively lipophilic molecules to diffuse into the deeper layer of the skin [3]. The thin layer (~20 μm) is composed of corneocytes and lipid matrix where latter one fills the extracellular space, and the whole assembly resemble a brick-and-mortar structure [4]. The passive permeation of external entity (molecules, particles, and formulations) is mainly controlled by the lipid matrix, and it is indeed responsible for barrier function of the skin [3,5]. A disturbance in the lipid composition and their lateral packing can lead to disease condition such atopic dermatitis and psoriasis [6].

A healthy skin barrier is necessary for defending the skin from the attack of foreign entities (pathogens, bacterial, and harmful substances), but at the same time it creates a hurdle in delivering the therapeutics on/through the skin [7,8]. Hence, several strategies are being developed and used to breach the skin barrier selectively and safely for numerous healthcare and personal care applications [9–11]. One of the strategies, is to apply external energy source (electric field [12], ion flux [13], and ultrasound [14]) or mechanical source such as microneedles [15] to name few. These methods require not only dedicated devices but also sophisticated control of operating parameters (electric field, voltages, pulse rate, microneedle size etc.). The other strategies focus on modifying the therapeutic formulation itself by incorporation of permeation enhancers [16], skin penetrating peptides [17], and nanomaterials [18].

The permeation enhancers are class of molecules known to enhance the skin permeation by either extraction of lipids or fluidization of skin matrix [19–21]. The penetration enhancers not only affect the lipid matrix but also the skin proteins which could lead to irritation and may have toxic effects [19,20]. The ethanol and dimethyl sulfide oxide are two common permeation enhancers, and it has been shown both by experiments [22,23] and simulations [24, 25] that they can disrupt the lipids packing to undesirable limits at higher concentrations.

Researchers have also studied nanoparticles (NPs)/nanomaterials as they have unique chemical and physical size dependent properties and can be tailor made from organic (lipids, biopolymers) and inorganic materials (metals) [26]. Based on the application one can choose either deformable nanomaterial like liposomes or rigid metallic nanoparticles. Liposomes are believed to cross the skin barrier by its deformation and merging with the skin lipid membrane, however exact mechanism are still unknown [27]. Gold NPs have been around since past few decades and been explored for various biological applications due to ease in modification of their surface by chemical or bioactive molecules to ensure the binding of different types of substrates over its surface [28]. The gold NPs of various shape, size and surface morphology have been synthesized and shown to be penetrating the deeper layer of the skin [29–33]. Not only experiments but also molecular simulations studies have been carried out to understand the permeation mechanism of NPs across the cell membrane [34,35] and skin lipid layer [36–37]. Molecular simulations provide a convenient way to understand permeation processes and can yield important physical insights with molecular detail that could not be obtained from experiments because of associated time and length scale. The in-silico models and methodology have been utilized for design of NPs of various morphology to overcome the barrier function of skin and intestinal membrane [38, 39] as well.

The simulation studies, so far presented in the literature, focus permeation of single nanoparticle across either the skin lipid or cell plasma bilayer [34–37]. However, several NPs are present in a formulation and when applied on skin, it is highly unlikely that nanoparticle-skin lipids interactions are not being affected by the other nanoparticle-nanoparticle interactions. There are some studies which have reported the co-operative transport of NPs across the cell membrane [40, 41]. Yue and Xianren Zhang reported the cooperative effect of nanoparticle permeation across the model lipid bilayer using the dissipative particle dynamics (DPD) simulations [40]. It was shown that small NPs formed small cluster on the membrane and internalized as a whole entity. Whereas the large nanoparticles remained dispersed and internalize independently. Four types of internalization pathways, namely, synchronous internalization, asynchronous internalization, pinocytosis-like internalization, and independent internalization were observed in simulations and found to be dependent upon the NPs size and their inter-particle distances. Zhang et al. performed DPD simulations of model membrane with NPs of various surface chemistry/patterns [41]. The translocation of NPs across the bilayer was observed in a cooperative manner and highly influenced with the surface chemistry of the beads of NPs. The hydrophilic NPs permeation increased with their concentration, but for partial and totally hydrophobic NPs, the opposite trend was reported.

Unlike cell membrane, skin lipid matrix is multilayer in nature and permeation mechanism could be very different from that of a single cell membrane. Moreover, in above studies [40, 41], the receptor mediated translocation process, was modelled by modifying the interaction parameters, whereas the transport across the skin SC layer is not receptor mediated. Hence, passive permeation of NPs of various size and surface chemistry across a multilayer model membrane would be entirely different. In this study, we report the translocation of NPs of various sizes, surface chemistry across the multilayer skin lipid matrix model. The unrestrained coarse grained (CG) molecular dynamics (MD) simulations were performed for bare, and thiol coated NPs. Two kind of bilayer systems, namely single lipid bilayer (SLB) membrane and double lipid bilayer (DLB) membrane were simulated, and it is shown that DLB membrane model should be used to study the translocation of nanoparticle across the skin lipid membrane.

## 2 Methods, Systems and Models

The permeation of molecules through skin lipid layers generally happens at milliseconds (ms) to micro-second (μs) time scale depending upon the size, shape, and surface chemistry of molecules. To capture the permeation events at realistic time and length scale, the CG models derived from MARTINI force field were used [42–44]. The NPs parameters were taken from earlier works [45, 46]. The CG structure of skin lipid bilayer (equilibrated for 3 μs) was taken from earlier work [39]. The bilayer size was 19.88 nm x 19.88 nm x 11.87 nm and had equimolar ratio of ceramide (CER), free fatty acid (FFA), and cholesterol (CHO). Here onwards this bilayer system is termed as single lipid bilayer (SLB) membrane model. Another model termed as double lipid bilayer (DLB) membrane model was prepared using the SLB. The water layer from the equilibrated SLB was removed, and remaining lipids were replicated in z direction and solvated again with water at both ends. The system was subjected to energy minimization, and subsequently NVT and NPT run [39].

To allow complete inclusion of nanoparticle in upper water layer and to avoid artifacts associated with the system size and nanoparticle–nanoparticle interactions over periodic boundaries, the water content was increased on each side of the bilayer. The NPs were inserted in the upper part of the water layer at the distance of ~6-7 nm from the center of the lipid bilayer. The overlapped water molecules were removed, and systems were energy minimized. The systems were further equilibrated for 200 ns each in NVT and NPT conditions by keeping lipids and NPs fixed to ensure the proper solvation. Properly solvated and equilibrated systems were then run for 1 μs in NPT ensemble.

All simulations were carried out in NVT and NPT ensemble using the GROMACS MD package [47–50]. The temperature was controlled at a skin temperature of ~305 K, using the Berendsen (equilibration run) and Nose-Hover (production run) thermostat with a time constant of 2 ps. Pressure was controlled by Berendsen (equilibration run) and Parrinello-Rahman (production run) barostat with a time constant of 6 and 12 ps respectively and compressibility of 4.5 x 10^-5^ bar^-1^ with semi-isotropic coupling. The pressure was independently controlled in XY and Z directions to obtain the tensionless lipid bilayer. The LJ potentials were smoothly shifted to zero between a distance rshift = 0.9 nm and the cutoff distance of 1.2 nm. The pair list was updated at every 20 steps. The configuration was sampled at every 100 ps in the production run.

## 3 Results and Discussion

### 3.1 Single lipid bilayer membrane model

The SLB models have been used to study the translocation of the nanoparticle across the cell and skin membranes [34–37]. To check, whether the same model can provide insights of permeation or translocation of multiple NPs, simulations with two types of NPs were performed. The bare nanoparticle had surface bead type of “C5” of MARTINI force field. The bare NPs coated with dodecane thiol were termed as coated NPs. The details of the systems simulated are given in the Table 1. The systems were labeled as ‘B’ and ‘C’ for bare and coated NPs respectively.

**Table 1.** Details of the simulated systems

| System | Membrane | Number of NPs | Size (nm) | Type | Simulation Time(μs) |
| --- | --- | --- | --- | --- | --- |
| B2 | SLM | 2 | 3 | Bare | 1 |
| B3 | SLM | 3 | 3 | Bare | 1 |
| B4 | SLM | 4 | 3 | Bare | 1 |
| B5 | SLM | 5 | 3 | Bare | 1 |
| B6 | SLM | 6 | 3 | Bare | 1 |
| B7 | SLM | 7 | 3 | Bare | 1 |
| B8 | SLM | 8 | 3 | Bare | 1 |
| B9 | SLM | 9 | 3 | Bare | 1 |
| C2 | SLM | 2 | 3 | Coated | 1 |
| C3 | SLM | 3 | 3 | Coated | 1 |
| C4 | SLM | 4 | 3 | Coated | 1 |
| C5 | SLM | 5 | 3 | Coated | 1 |
| C6 | SLM | 6 | 3 | Coated | 1 |
| C7 | SLM | 7 | 3 | Coated | 1 |
| C8 | SLM | 8 | 3 | Coated | 1 |
| C9 | SLM | 9 | 3 | Coated | 1 |
| B13 | DLM | 3 | 1 | Bare | 1+3* |
| B16 | DLM | 6 | 1 | Bare | 1+3* |
| B19 | DLM | 9 | 1 | Bare | 1+3* |
| B33 | DLM | 3 | 3 | Bare | 1+3* |
| B36 | DLM | 6 | 3 | Bare | 1+3* |
| C13 | DLM | 3 | 1 | Coated | 1+3* |
| C16 | DLM | 6 | 1 | Coated | 1+3* |
| C19 | DLM | 9 | 1 | Coated | 1+3* |
| C33 | DLM | 3 | 3 | Coated | 1+3* |
| C36 | DLM | 6 | 3 | Coated | 1+3* |

\*Extended runs

NPs of diameter 3 nm were utilized for studying the interaction with SLB model. MD simulations were performed by initially placing the NPs at the top of the SC membrane (Figs. 1A, and 1B) as described in section 2. As summarized in Table 1, simulations were performed for 1 μs with 2, 3, 4, 5, 6, 7, 8, and 9 NPs. The snapshots of systems at the end of the simulation run are shown in the Fig S1 (see supporting information). Spontaneous partitioning of NPs was observed in all simulations (Figs. 1, S1, and S2), irrespective of the surface coating and concentrations. NPs have exhibited events of partitioning as individual monomers (Figs. 1A and S1) as well as for the aggregates (Figs. 1B and S1). These findings highlight a highly favorable environment offered by the SC membrane for the NPs. Membrane configurations indicate alteration in the membrane packing after the partitioning of the NPs, and the distortion appears to be more dominating for the coated NPs (Fig. S1).

**Figure 1.**
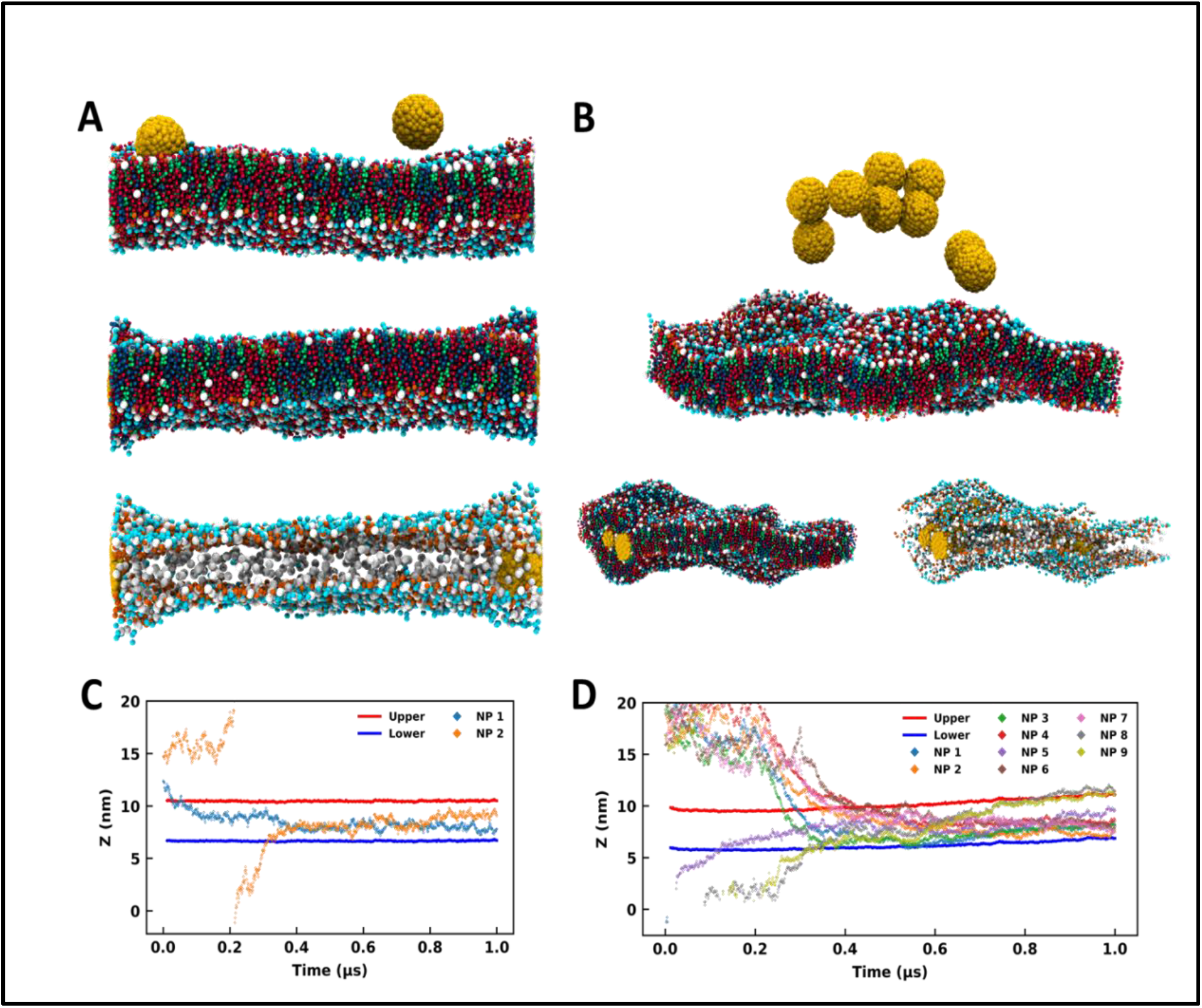
Bare and coated nanoparticles partitioning into the SLB model. Side-view MD snapshots of the (A) B2 and (B) B9 systems having initial (top) and final configurations with and without lipid molecules. Time trace for z-coordinates of the center of mass of NPs, and CER headgroups of upper and lower leaflets for (C) B2 and (D) B9. The CER, FFA, CHO. NP are represented in red, blue, green, and yellow color respectively. The images were rendered using VMD software [58].

The above results are in line with some of the experimental evidence of finding the gold NPs in the deeper layer of the skin [29–33]. Huang et al. showed that gold NPs could deliver the model protein drugs in the skin dermis [29]. The finding of hydrophobic NPs in the interior of the lipid bilayer have been reported in some of the earlier simulation studies of plasma membrane as well [35, 37, 51]. The fullerene like NPs (size ~1.2 nm) also have been found in the interior of the phospholipid bilayer and it’s been shown that the NPs remained dispersed in the interior of the bilayer even at moderate concentrations of NPs [34, 52–55]. The bare nanoparticle (3 nm) and coated (~ 5 nm) are bigger than the bilayer width and found to be agglomerated even at moderate concentrations (Fig. S1). This implies that not only concentration of NPs but also NPs size determine their organization in the membrane environment.

We have analyzed trajectories of center of mass of ceramide headgroups (upper and lower) and center of mass of each NPs along the z-direction to investigate the partitioning behavior of the NPs (Figs. 1C, 1D, and S2). For the B2 system with two bare NPs (Fig. 1A), one NP was initially placed at the membrane-water interface while the other was placed above the SC membrane and in the bulk water (Figs 1A and 1C). Spontaneous insertion was observed for both the particles (Figs. 1A and 1C), while the first particle got partitioned within initial 100 ns, the other one in bulk water entered the membrane after 300 ns. Once inserted, the NPs were present in the membrane core for the rest of the trajectory. The swelling of the membrane in the vicinity of NPs is quite apparent from the configurations (Fig. 1A) due to the hydrophobic mismatch between the membrane thickness and the size of the particles.

For the B9 system, the higher concentration of the NPs resulted in the aggregation in bulk water (Fig. 1B). Interestingly, the membrane allows the partitioning of this aggregate and accommodates a high concentration of NPs (Fig. 1B). Although membrane packing appears to be significantly distorted due to the lipid reorganization in the vicinity of the NPs, the overall bilayer structure is sustained. From the trajectory time trace data (Fig. 1D and Fig. S2), it can be observed that five NPs were part of the aggregate, which is evident from the strongly correlated z-coordinates of those particles and insertion from the upper leaflet. However, one NP got partitioned into the membrane within initial 100 ns from the downside. Two NPs formed another aggregate that entered the membrane environment from the lower leaflet. Although both aggregates entered the membrane at quite close time points, the membrane accommodated all particles while maintaining the bilayer structure (Fig. 1B).

All other systems (except B3 and B8) with bare and coated NPs exhibited similar behavior (Fig. S1) where the NPs partitioned into the SC membrane within 500 ns (Fig. S2). B3 system presented an exception where the particles formed an aggregate in the initial stage and got partitioned into the membrane towards the end of the 1 μs simulation (Fig. S2). Nevertheless, the partition was witnessed in all the simulations. In B8, the distortion is quite significant, as apparent from the membrane configuration (Fig. S1). Higher distortion in the membrane topology in the presence of coated particles compared to the bare particles (Fig. S1) is indicative from the time trace data for headgroups which shows significant displacement in the membrane trajectories for coated particles.

The bilayer structure of the SC membrane has been altered due to the interactions with the NPs. The reassembling of lipids and distortion in the membrane is apparent from the membrane configuration corresponding to the last frame of 1 μs simulations (Figs. 1B and Fig. S1). To quantify the perturbation in the membrane system due to the presence of higher number of NPs, the head group fluctuations were evaluated. We have calculated distributions of the CER headgroup fluctuations for the entire 1 μs trajectory and the last 100 ns of the trajectory to comprehend the alteration in membrane topology (Figs. 2 and Fig. S3). It is evident from the headgroup distribution for 1 μs data that the coated NPs have significantly perturbed membrane headgroups (Fig. S1). The membrane has shown significant displacement for C5 as compared to the bare counterpart B5 (Figs. 2A and S2). Also, the spread for the coated particles (~5 nm) is significantly larger than the bare particle (~3 nm). Higher fluctuations in the presence of coated particles compared to the bare particles are intuitive due to the presence of protruding thiol groups in the former.

**Fig. 2.**
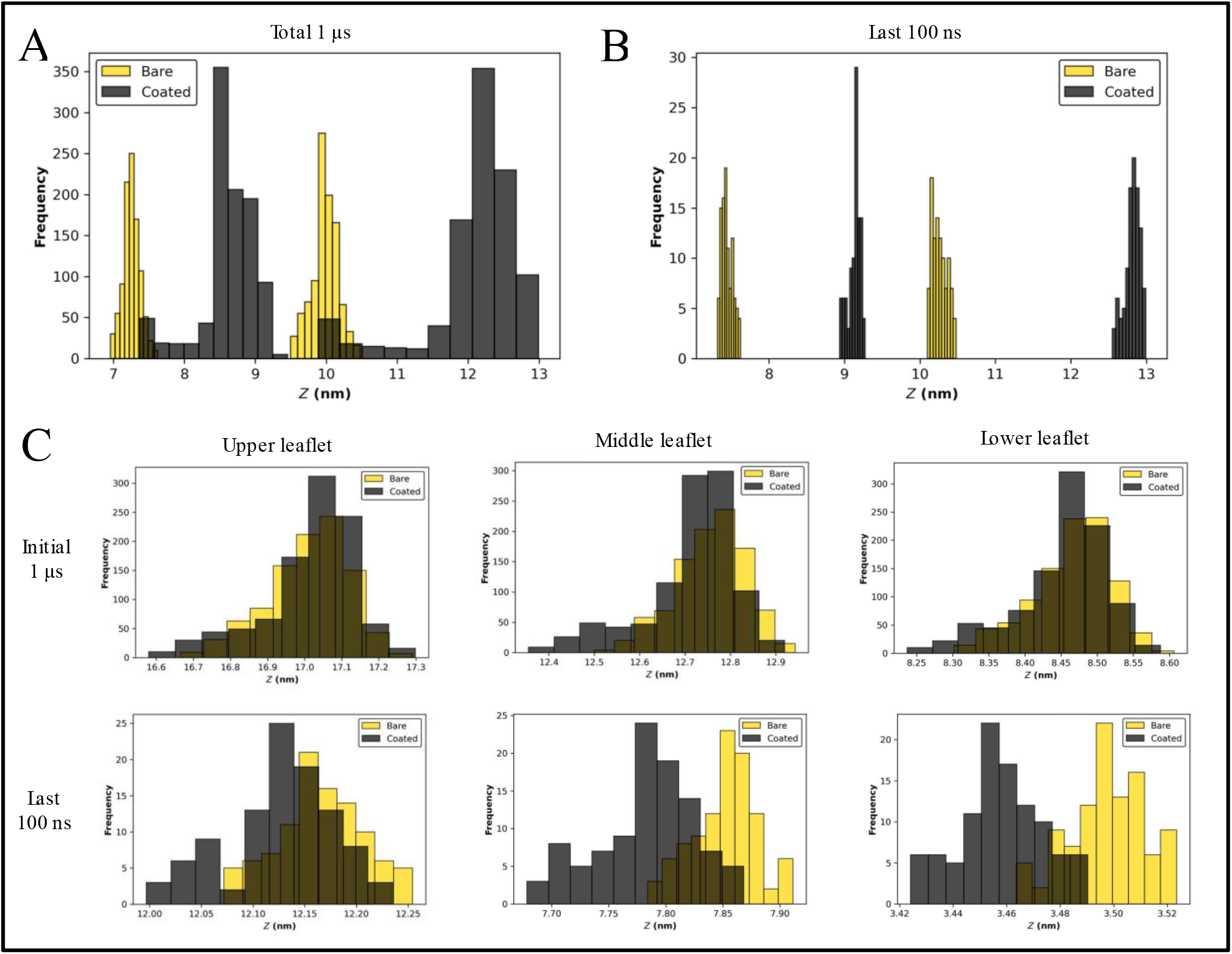
SLB model exhibit unrealistic headgroup fluctuations. Distribution for CER headgroup fluctuations for B5 (gold) and C5 (black) for the (A) entire trajectory and (B) last 100 ns. (C) Distribution for CER headgroup fluctuations for B13 (gold) and C13 (black) for the entire trajectory (top) and last 100 ns (bottom).

Final configurations suggest a stable membrane for both bare and coated NPs (Fig. S1). Consequently, we have also determined fluctuations for the last 100 ns trajectory (Figs. 2B and Fig. S3). Interestingly, it can be perceived that the fluctuations are settled towards the end of simulations for both cases, highlighting a tendency of the membrane to accommodate both types of NPs. The fluctuations are of the same order for both bare as well as coated. During insertion, it can be concluded that perturbation in membrane headgroups for coated NPs is significant compared with the bare NPs (Figs. 2A and S3). Although once the membrane gets equilibrated after accommodating the particles, the fluctuations become similar for both particles (Figs. 2B and S3). These trends of higher distortion in the presence of coated NPs are conserved for all the cases (Figs. S1 and S3), except for B8, where the membrane is entirely distorted (Fig. S1), and hence a head-on comparison is not feasible. Also, all the simulations manifest undulated membranes towards the end of trajectories (last 100 ns).

The presence of NPs undulated the membrane as can be seen from the projected area of bilayer on XY plane (Fig. S4). Irrespective of nanoparticle type (bare or coated) the area decreased with increase in the number of NPs, however for any given number of NPs, the projected area was smaller for coated nanoparticle as compared to the bare one. This implies that coated NPs induced more undulation in bilayer as compared to the bare NPs. The stability of the bilayer can also be depicted from the density profile shown in Fig. S5. The presence of dual peaks in the lipid’s density corresponds to the existence of bilayer structure. The dual peaks got disappeared with increased number of NPs for both bare and coated case. The swelling of the bilayer is also apparent from the increased spread of density in z direction. The tail order parameter (Fig. S6) again confirms that, presence of higher number of NPs significantly affected the lipid packing.

The order parameter decreased with the increased nanoparticle numbers and coated nanoparticle had much pronounced effect on order parameter as compared to bare for a given concentration of NPs. The bare NPs created small clusters in the interior of the membrane, whereas coated NPs created a single large cluster (Fig. S7-Fig. S10). Hence, membrane with coated NPs were found to be in highly undulated form.

The SC membrane comprises three distinct types of lipids: CER, FFA, and CHO. To understand if translocation of the NPs with the SC membrane is being stimulated by any selective lipid or all of them, we have examined their interactions independently. We have perceived all three lipids in the proximity of the NPs (Fig. 3A), which highlights strong interactions of the NPs with all three lipids. To understand the correlation within all components, we have evaluated the lateral density of all four components: NPs, CER, CHO, and FFA. The high lateral density of CHO molecules in the vicinity of the NPs exhibits a strong tendency of CHO molecules to bind with the NPs (Figs. 3B and Fig. S7-S10). Although this correlation contrasts in the case of the CHO molecule it also persists for CER and FFA (Figs. S7-S10). Interestingly these trends appear to be consistent for both types of particles (Figs. S7-S10).

**Fig. 3.**
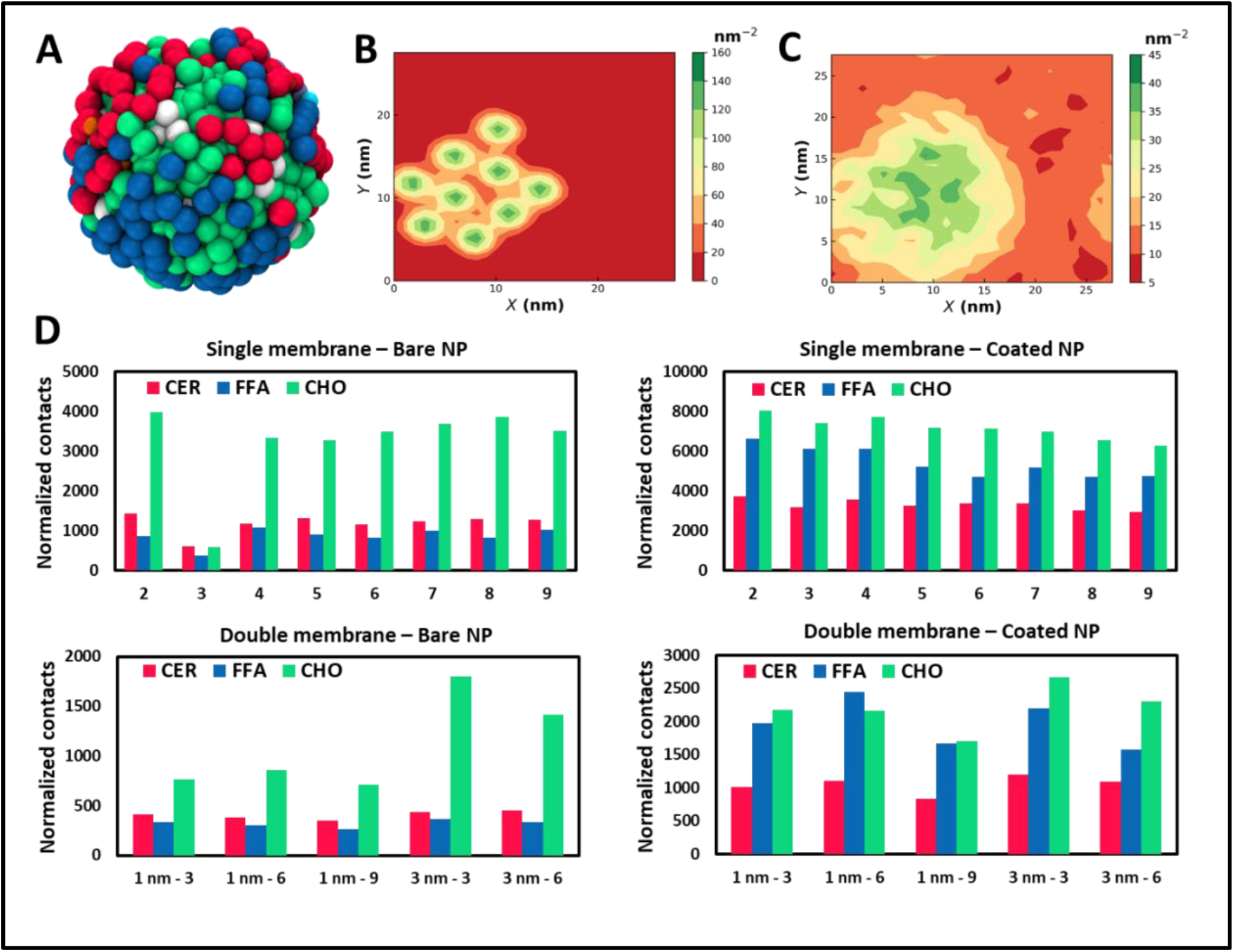
Lipid molecules show selective binding with NPs. (A) MD snapshot of the lipid molecule within 1 nm of a gold NP. Lateral density map of (B) NPs and (C) CHO for B9. (D) Normalized contacts of lipid species with the NPs. Contacts are calculated for the last 100 ns of trajectories and normalized with the number of MARTINI beads corresponding to each lipid species and the number of NPs in the system. The CER, FFA and CHO are represented with red, blue, and green color respectively.

To quantify these interactions and have a more robust gauge for comparison between lipid species, we have performed contact analysis over the last 100 ns of trajectory. We have calculated total contacts made by NPs with the lipid species, where contact is defined when a lipid CG bead is present within the radius of 1 nm of an NP. The normalized contacts (Fig. 3D) are calculated by normalizing the total contacts over trajectory by the number of NPs in each system and the number of beads present in individual lipid species. A strong interaction of NPs with all three components can be recognized, and hence the NPs partition into the membrane. We observe a preferential binding of CHO molecules compared to CER and FFA in all the cases (Fig. 3D). The strong interaction of NPs with CHO highlights the capability of CHO as an effective receptor for NPs. For systems corresponding to bare NPs, the differences between CHO and the other two species are remarkably distinct. Although B3 shows considerably lower contact due to the slow permeation process than the other cases (Fig. S2). It can also be observed that the bare NPs have a stronger tendency to interact with CER when compared with the FFA. The trends for normalized contacts are quite reproducible across all the cases showing a strong preference of CHO binding with the NPs irrespective of concentration and aggregation.

For coated NPS, there is a notable difference in trends of contact analysis. The total contacts are on the higher end for coated particles than the bare particles due to the protruding thiol groups, which provide additional interaction site. Particularly relative intensification in FFA contacts can easily be recognized. The tendency of preferential interactions of bare NP with the CER compared to the FFA gets interchanged in the case of coated NPs, where the FFA shows an enhanced binding in the vicinity of the NPs. Indeed, CER and FFA contacts are comparable to the CHO contacts for coated NP, while there was a significant difference in the case of bare NP. The finding of the NPs in the vicinity of the CHO can also be seen from the radial distribution function (Fig. S11) plots.

### 3.2 Double lipid bilayer membrane model

In section 3.1, the results of interaction of NPs of various size and surface chemistry with SLB model at various concentration of NPs are discussed. High concentration of NPs induced several structural changes in the bilayer and one reason could be due to presence of water layer at both ends of the bilayer. In the skin SC, several lipid layers are arranged in lamella form.

A SLB model for the SC membrane is deficient in representing the multilayer architecture of this membrane. To ascertain whether the trends addressed hitherto are realistic and not an artifact due to the simplistic SLB model, we have employed a multilayer (two layers) model to represent the skin barrier, which is closer to real system. Also, to understand the effects of NP size on the partitioning properties, we have studied two sizes of NPs (1 nm and 3 nm) using this DLB (Figs. 4A and 4B). We have studied three systems for 1 nm particles with three, six, and nine NPs (Table 1 and Fig. S12). For 3 nm particles, two simulations were performed using three and six NPs (Table 1 and Fig. S12). Two bilayer simulations were carried out in a stepwise manner, where in the first step NP partitioning was observed for a duration of 1 μs. The water molecules were reduced in the second step to reduce the system size, and simulations were extended for another 3 μs. The assertive partitioning behavior observed in the SLB model was effectively reproduced in the multilayer model (Figs. 4 and S12). Like what was discerned in the SLB model, the NPs were partitioned into the DLB SC membrane within the initial 600 ns of simulations, except for C33 (Fig. S13 and Fig. S14).

**Figure 4.**
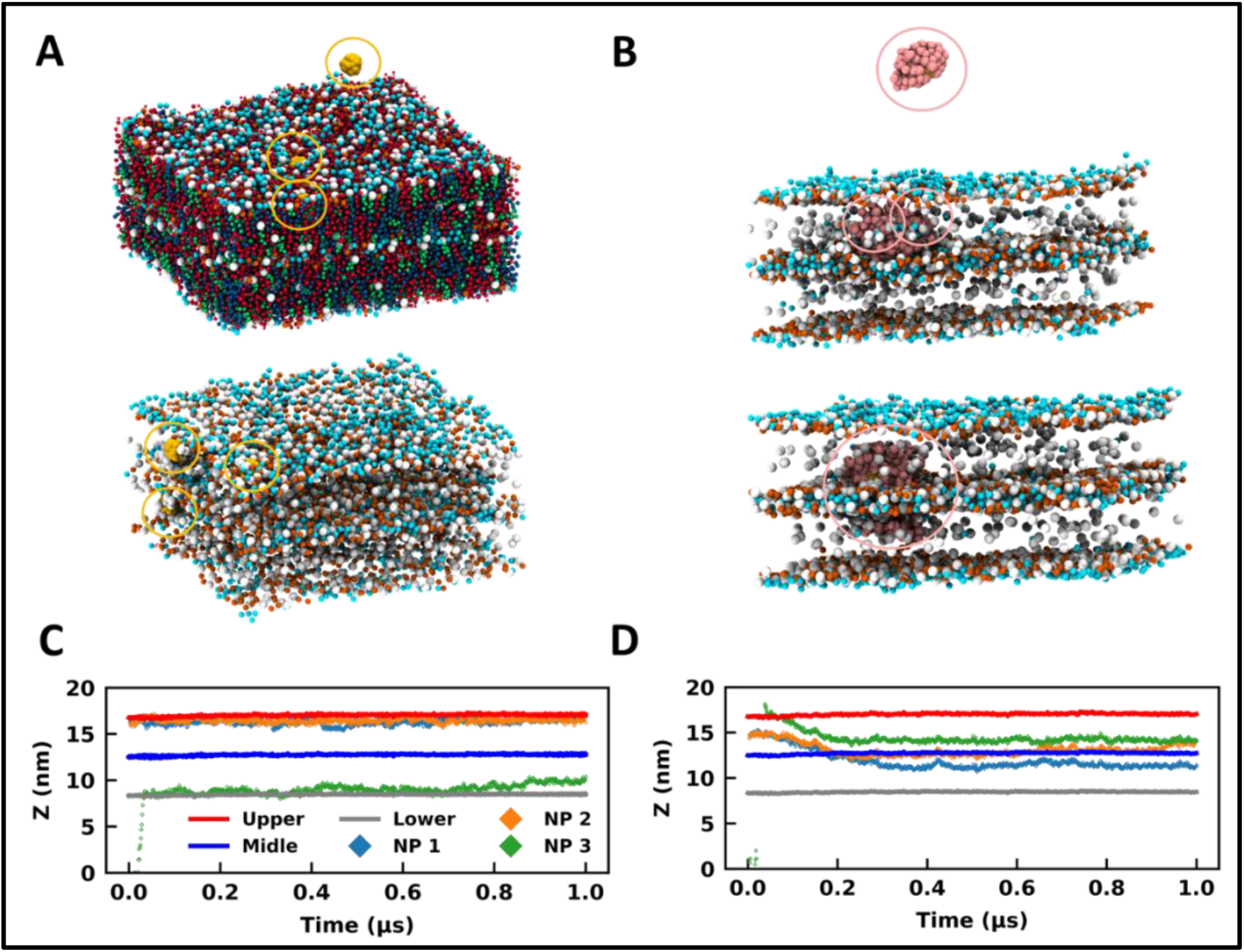
Bare and coated nanoparticles partition into the DLB model. Side-view MD snapshots of the (A) B13 and (B) C13 system having NPs and a double-bilayer SC model corresponding to the initial (top) and final (bottom) configuration. Lipids are hidden in some cases for clarity. Time trace for z-coordinates of the centre of mass of NPs, and CER headgroups of the upper, middle, and lower leaflets for (C) B13 and (D) C13. Same colour scheme used for both plots. The CER, FFA, CHO. NP are represented in red, blue, green, and yellow colour respectively. The NPs are highlighted in circles. The images were rendered using VMD software [58].

In B13, two NPs were initially placed in the headgroup region of the membrane, and the third one was placed in bulk water above the DLB model (Fig. 4A). All three particles were restricted to the headgroup region of the membranes for the entire trajectory (Figs. 4A and 4C). It can be concluded from the trajectories of the 1 nm bare particles in all simulations that a favorable site is present for such molecules in the headgroup region of the membrane (Figs. 4A, 4C, and S13). The membrane headgroup region provides a favorable environment for the 1 nm particles even at high concentrations (Fig. S13). Similar behavior can also be observed for the 3 nm size bare particle where the NPs prefer sampling the headgroup regions in the initial part of trajectories (Fig. S14). Although later on the NPs were present in the membrane core. Irrespective of the concentrations, the trend is adequately reproduced across systems. The NPs demonstrate characteristics considerably like partially amphiphilic small molecules, which prefer to sample the headgroup region of the lipid membranes [57]. It appears that the bare particles remained confined to the membrane (top or bottom) they initially partitioned through (Figs. S13 and Fig. S14). The preference for this confinement was witnessed as headgroup region for 1 nm particles and membrane core for 3 nm particles. However, if we observe the trajectories for the additional 3 μs, we can observe a few events where the particles have reached the interface of two bilayers for B19 and B33 (Figs. S15 and S16). Hence, transport across the multilayer skin barrier for these particles can be inferred as a rare event. As discussed earlier, the gold NPs can easily cross the skin barrier and found to be in the deeper layer. To reach into the deeper layer of the skin, the NPs must cross several layers of the lipid layers.

For the coated particles, C13 was created by initially placing two NPs in the membrane core of the top bilayer (Fig. 4B). One particle was placed in bulk water above the multilayer model (Fig. 4B). The coated NPs have distinct interactions with the membrane when compared with the bare NP (Figs. 4B and 4D). It can be observed that the thiol coated particles have a preference to sample the interface of two bilayers (Figs. 4B and 4D), which can be observed from the trajectories of particles along the z-direction (Fig. 4D). This behavior is independent of the particle size and is found for both 1 nm and 3 nm particles (Fig. S13-S16). Though it is a strong function of the surface coating and hence was rarely observed for the bare particle as discussed previously. It is vital to sample the interface between two membranes for effective transport across membranes. Coated NPs have exhibited multiple events of sampling across the membranes, highlighting their capability to cross the multilayer skin barrier. So far, we have observed the spontaneity of gold NPs to partition into the SC membrane and how these trends get altered due to the surface characteristics of the NP. The necessity to utilize a multilayer model is noteworthy concerning its capability to study the transport across membranes, which the SLB model disregards.

The NPs cross the skin barrier via passive diffusion. Earlier studies by Yue et al. [40] and Zhang et al. [41] showed that in case of receptor mediated transport, small NPs formed clusters and translocated inside the membrane whereas bigger sized nanoparticle penetrated independently. The hydrophilic NPs translocation increased with increased in their concentration but reduced for partial and totally hydrophobic NPs. In our simulations, we haven’t observed any such specific translocation events, however, irrespective of membrane models, nanoparticle types, the NPs were found in the membrane interior. The coated nanoparticle formed bigger cluster inside the membrane interior as compared to the bare NPs.

We have witnessed high headgroup fluctuations in the SLB model of the SC in the presence of NPs. Intending to determine the dependency of these fluctuations on the membrane model, we have performed a similar analysis for the DLB model. The comparison of headgroup fluctuation data between B13 and C13 for the initial 1 μs trajectory explicate an insignificant (~0.1 nm) difference between the distributions for bare and coated NPs (Fig. 2C). Fluctuation data for the last 100 ns of the 3 μs trajectory also exhibit differences of the same order between bare and coated (Fig. 2C). The fluctuation data for all the DLB demonstrate a similar trend (Fig. S17). The lipid density (Fig. S18) profile also confirms that even at higher concentrations of the NPs the bilayer holds its structure and undulations were less as compared to the SLB. A similar conclusion can also be made based on the order parameter data for the DLB (Fig. S19), which shows lesser disorder in the DLB than the SLB. Though to a lesser extent, the order parameter calculations show a higher disorder in coated particles than the bare. With these findings, we can conclude that the DLB model does not manifest high fluctuations like what was witnessed for the SLB. A multilayer or a DLB model should be employed to model the skin barrier.

Lateral density maps show a correlation between the NP density and the CHOL density for the DLB (Figs. S20-S22), like the SLB (Figs. S4-S7). However, the correlation is not as assertive as it was for the SLB. Dissenting to the SLB model’s findings, the DLB model’s density maps indicate an opposing trend. The lateral density for CER and FFA appears to be lower at the sites corresponding to the NPs. Interestingly this is observable even when the densities for lipid species are higher than the SLB.

Analogous to the SLB, we have calculated contacts in this model for all cases. However, there is a considerable decrease in the value of normalized contacts for the DLB compared to the single counterpart. The reduction in the case of CHO is more significant when compared with the other two species. The radial distribution function plots (Fig S23 and Fig. S24) also confirm these findings. Nevertheless, the relative trends are conserved from the SLB model. CHO molecules show preferential binding as compared to CER and FFA. This difference gets significantly reduced for coated NPs. Furthermore, opposite trends for preferential binding for CER and FFA in bare and coated NPs are reproduced as they were in SLB. The results highlight the selectivity of lipid species to bind to NPs based on the surface chemistry preferentially, and it is conserved in both models.

## Conclusion

NPs are being utilized for several personal care and healthcare applications. We have employed coarse-grained MD simulation to elucidate the interactions of bare and surface coated gold NPs with the single-bilayer and double-bilayer model of the Stratum Corneum. Our study highlights the differences that arise in the permeation characteristics of the NPs due to the surface coating. The NPs exhibit a strong tendency to partition into the SC membrane, irrespective of their surface coating. Both SLB and DLB model provides a highly favorable environment for NPs. The SLB model gets significantly destabilized in the presence of coated NPs compared to the bare NPs as exhibited by the head group fluctuations, order parameter, and density distribution calculations. The coating of thiol groups is found to overcome the incapability of bare NPs to translocate across the membranes in the DLB. Although, the fluctuations in the SLB seem unrealistic because they are not present in the DLB, which is a more accurate replica of the skin barrier. Our contact analyses highlight the strong tendency of lipid components of the SC layer to interact with the NPs, albeit CHO interacted more as compared to FFA and CER. Also, the interactions of FFA and CER with the NPs are significantly different across the bare and coated NPs.

These findings would certainly play a prominent role in targeted drug delivery across the skin barrier. The NPs insertion into the SC membrane is found to be a vital function of the size and surface chemistry of the NPs. Also, targeted delivery can be employed based on the preferential interactions of the NPs with the lipid species. Our study emphasizes significant characteristics of NP-SC interactions, which highlight the necessity of effective NPs synthesis for transport through the skin barriers. It also highlights the necessity of employing more realistic multi-layer models for studying the barrier offered by the SC.

## Abbreviations

NPs: Nanoparticles
SC: Stratum corneum
MD: Molecular dynamics
CHO: Cholesterol
FFA: Free fatty acid
CER: Ceramides
SLB: Single lipid bilayer
DLB: Double lipid bilayer
DPD: Dissipative particle dynamics
CG: Coarse-grained

## Authors contribution

R.G. and B.R. conceptualized the idea. Y.B., P.S. and R.G. contributed equally. All authors contributed to discussing the results and writing, editing the manuscript.

## Acknowledgement

Authors would like to thank, Dr Gautam Shroff, Head, TCS Research and Mr. K Ananth Krishnan, CTO and EVP, Tata Consultancy Services for their constant encouragement and support during this project.

## Funding & Resources

This research was funded by Tata Consultancy Services (TCS), CTO organization.

## Conflict of interest

The authors declare no conflicts of interest.

## Notes

### Competing Interest Statement

The authors have declared no competing interest.

