## Supporting Information for "Elucidating Collective Translocation of Nanoparticles Across the Skin Lipid Barrier: A Molecular Dynamics Study"

‡contributed equally

### Supporting information

### 1. Single bilayer

#### 1.1 Snapshots

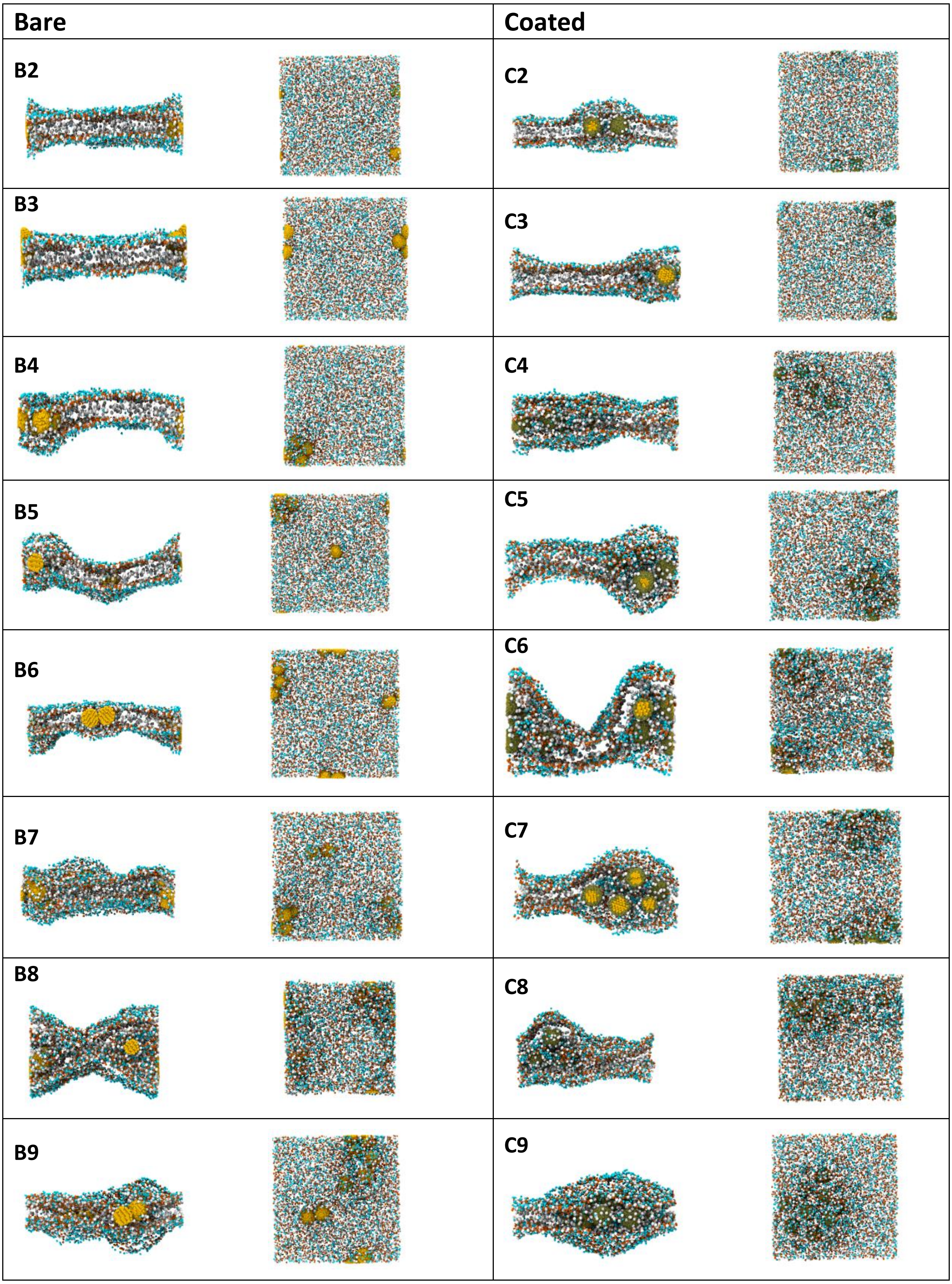


**Figure S1.** Final snapshots of single bilayer-NP system after 1 μs simulation run.

#### 1.2 Trajectories

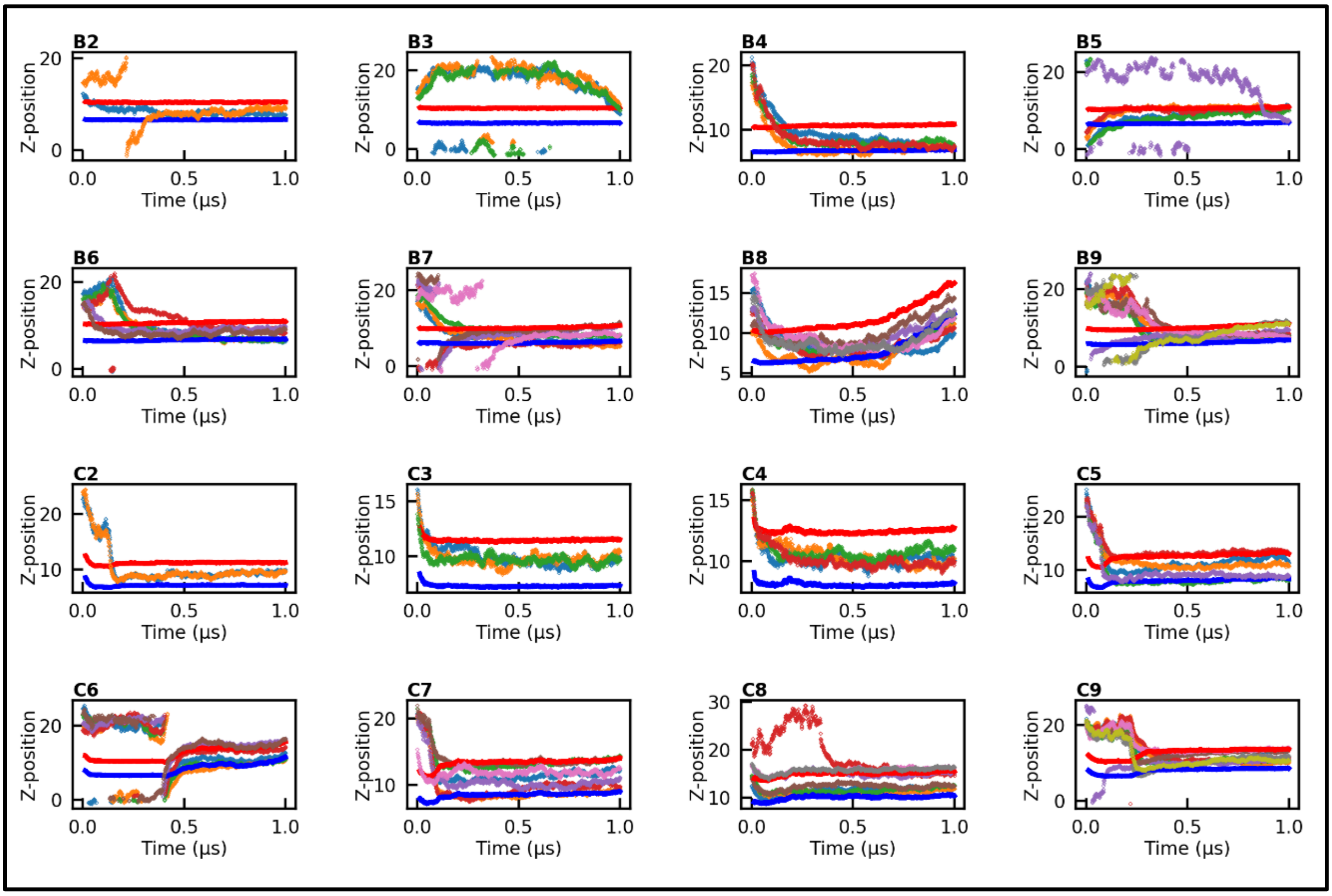


**Figure S2**. Time trajectories of bilayer headgroup atoms and NPs along the z direction calculated over 1 μs simulation time. The head group data for upper and lower leaflet are shown in red and blue color respectively. Individual NP trajectories are shown in other colors.

#### 1.3 Fluctuations

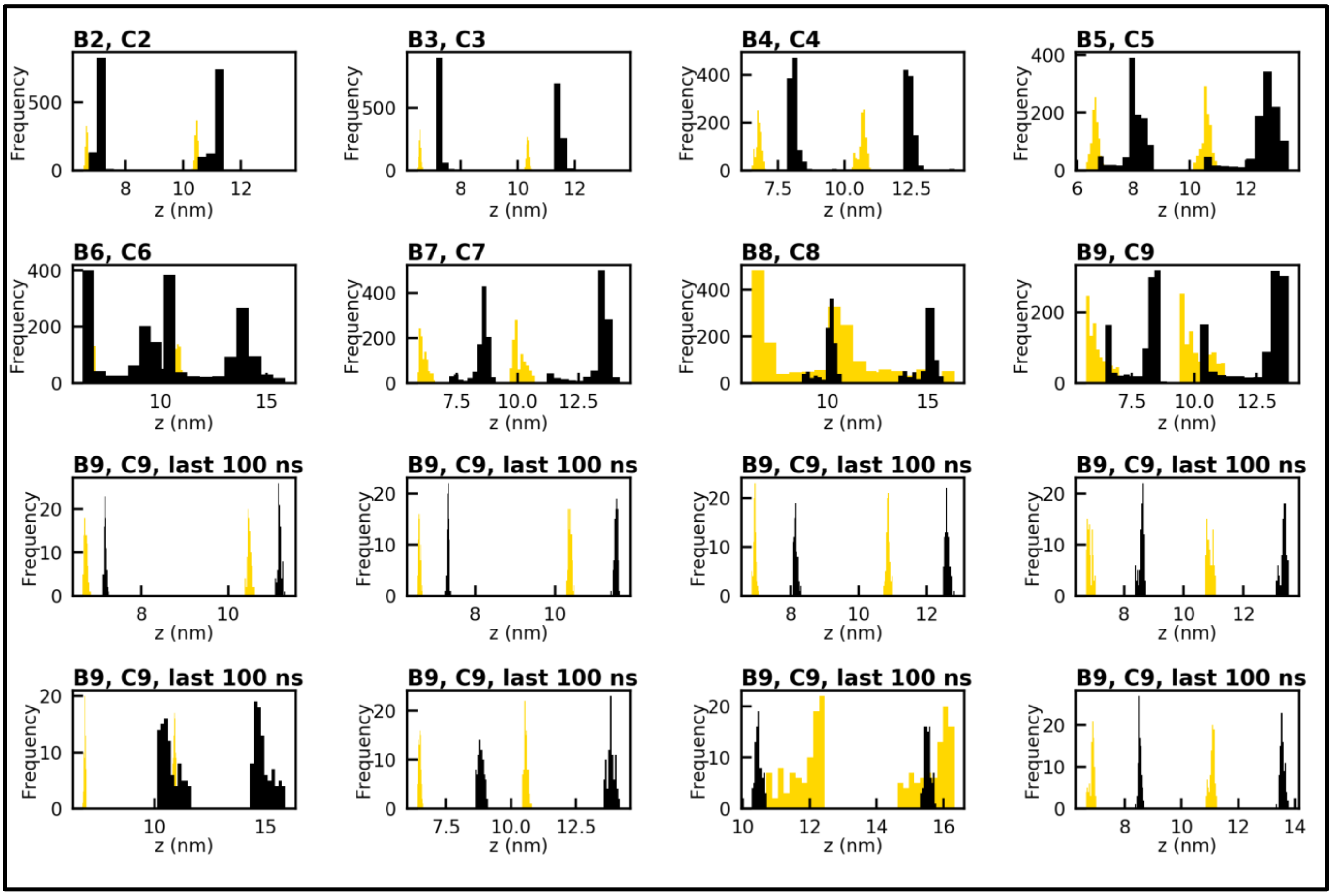


**Figure S3**. Histograms of time trajectories given in Figure S2. The narrower histograms for last 100 ns trajectory signify stable bilayers as compared to entire 1 μs trajectory. Bare and coated NPs are shown in gold and black color respectively.

#### 1.4 Projected are on XY Plane

The projected area on xy plane per lipid calculated using the following equation:

$$A = \frac{c\,Lx\,Ly}{Nlipid} \qquad \text{(S1)}$$

Where $L_x$, $L_y$ is the box length in X and Y direction, respectively, the C takes value of two and four for single a double bilayer respectively, and $N_{lipid}$ is total number of lipids in the bilayer.

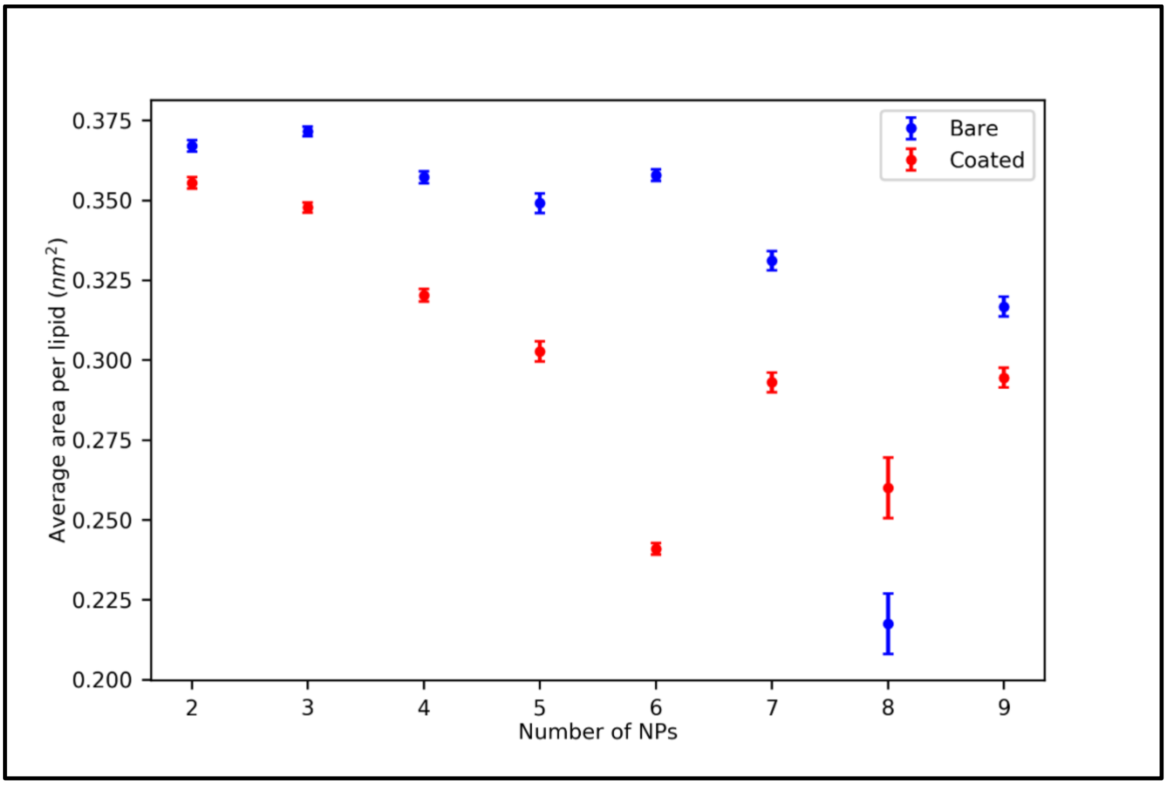


**Figure S4.** Projected area on XY plane of bilayer calculated in last 0.1 µs production run.

#### 1.5 Bilayer density along bilayer normal (z axis)

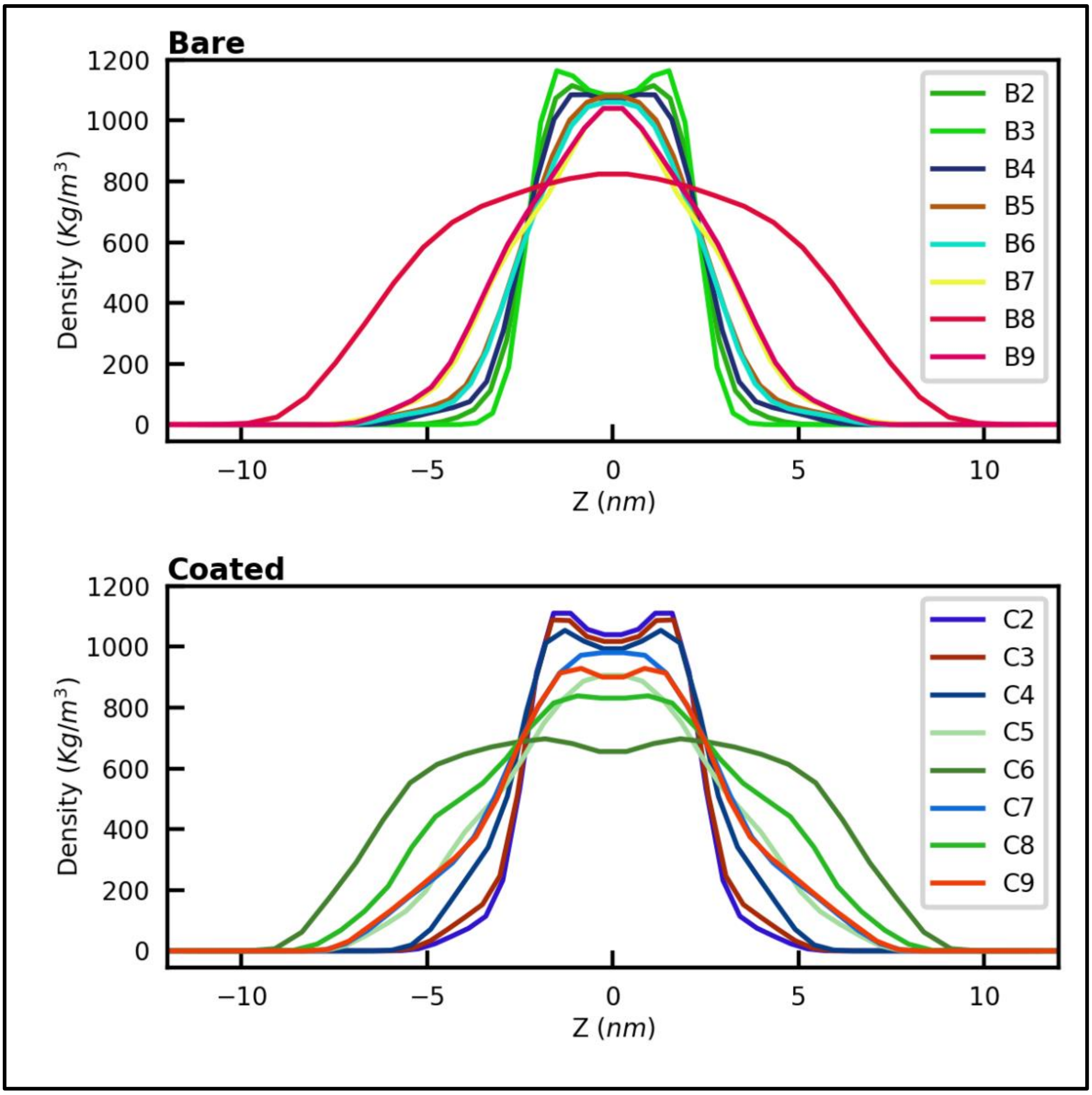


**Figure S5.** Bilayer density along z axis (last 100 ns)

#### 1.6 Lipid order parameter

The second rank order parameter for the bilayer, which has normal in z direction, could be defined as:

$$Sz = \frac{1}{2}(3cos^2\theta - 1) \qquad \text{(S2)}$$

where θ is the angle between the bond and the bilayer normal. Sz = 1 means perfect alignment with the bilayer normal, Sz = −0.5 anti-alignment, and Sz = 0 random orientation of the lipid chains.

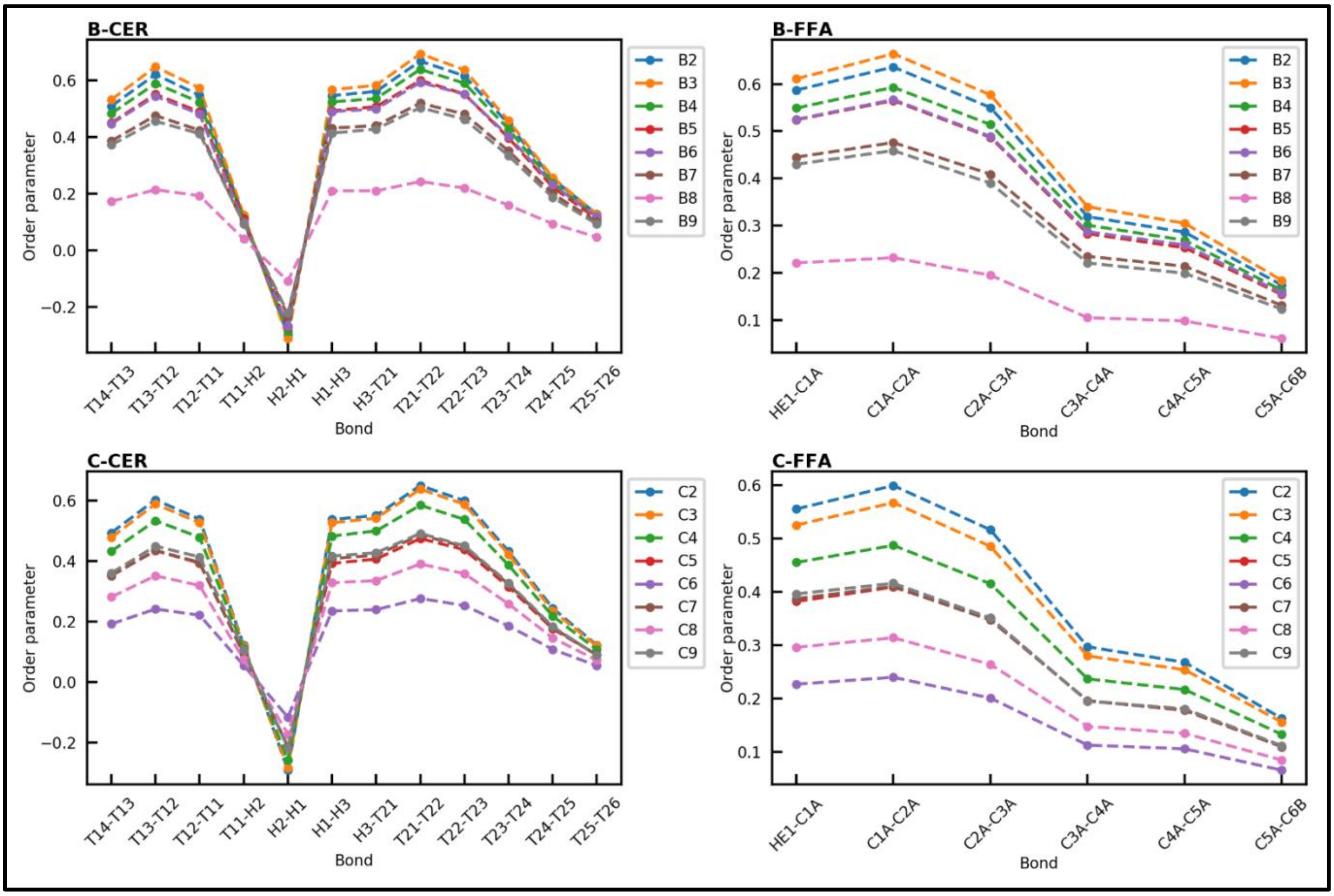


**Figure S6.** Tail order parameter of CER and FFA chains. B and C corresponds to bare and coated nanoparticle system respectively.

### 1.7 Density maps

#### 1.7.1 Bare

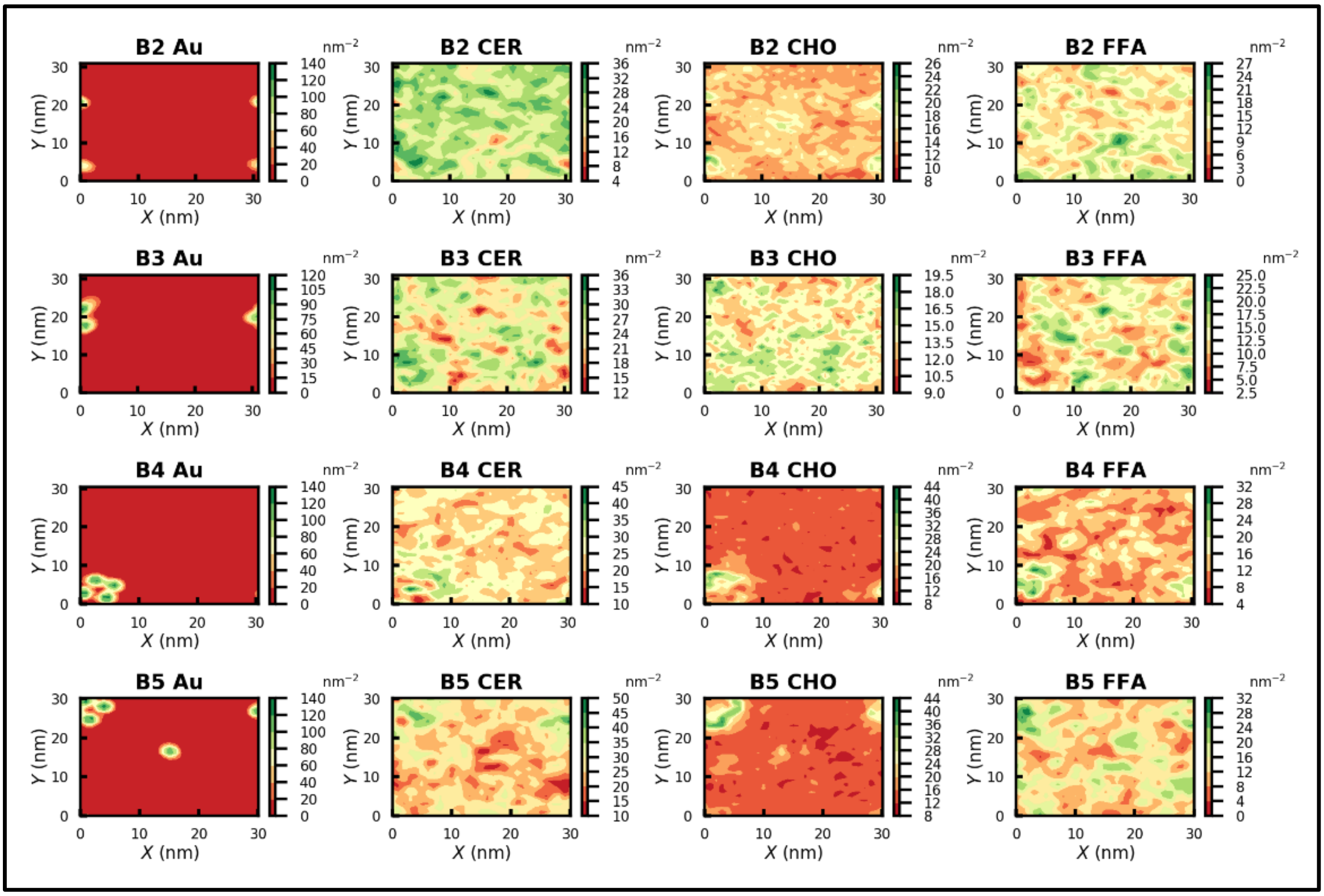


**Figure S7.** Density maps of nanoparticles, ceramide, cholesterol, and free fatty acid for B2 to B5 systems.

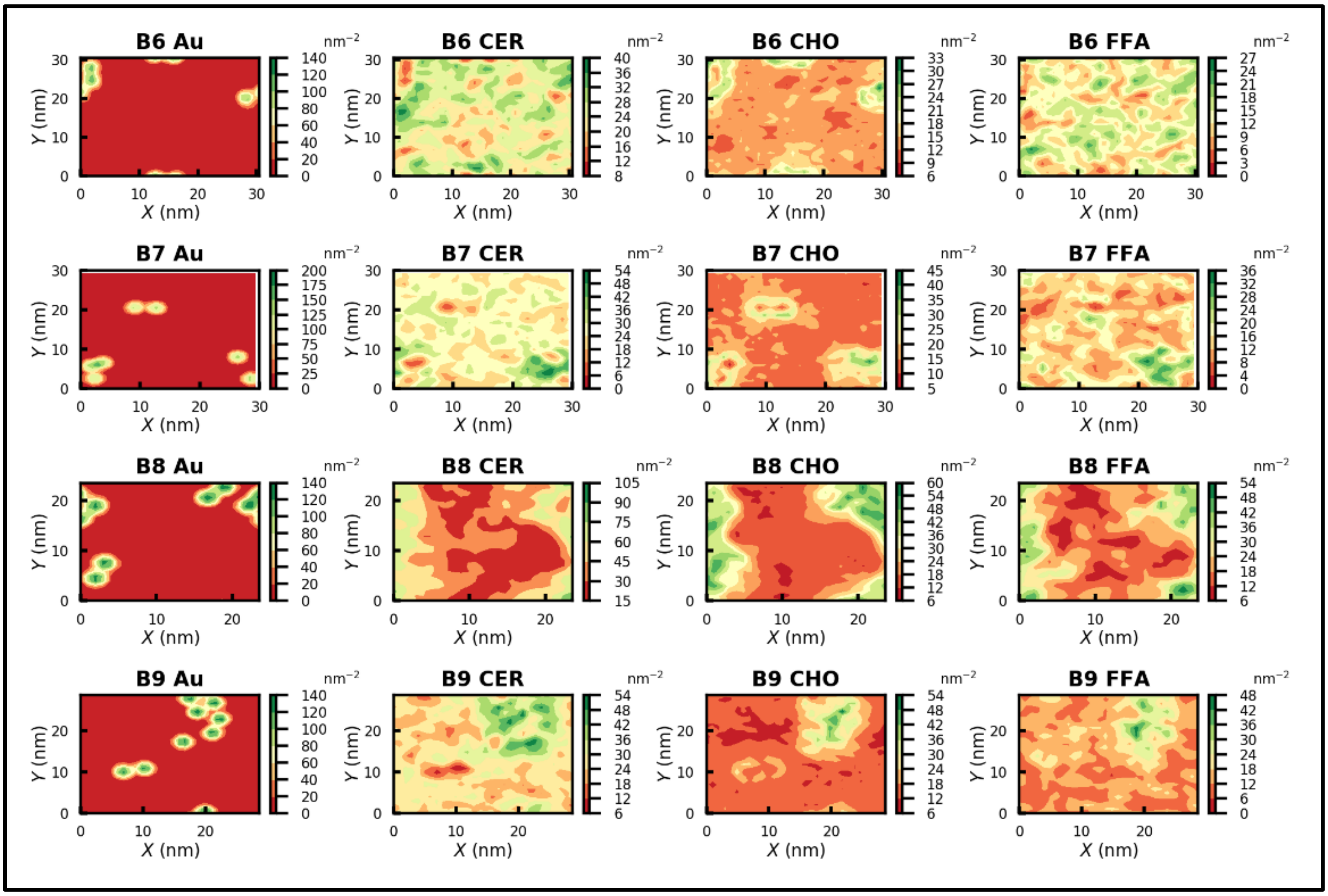


**Figure S8.** Density maps of nanoparticles, ceramide, cholesterol, and free fatty acid for B6 to B9 systems.

#### 1.7.2 Coated

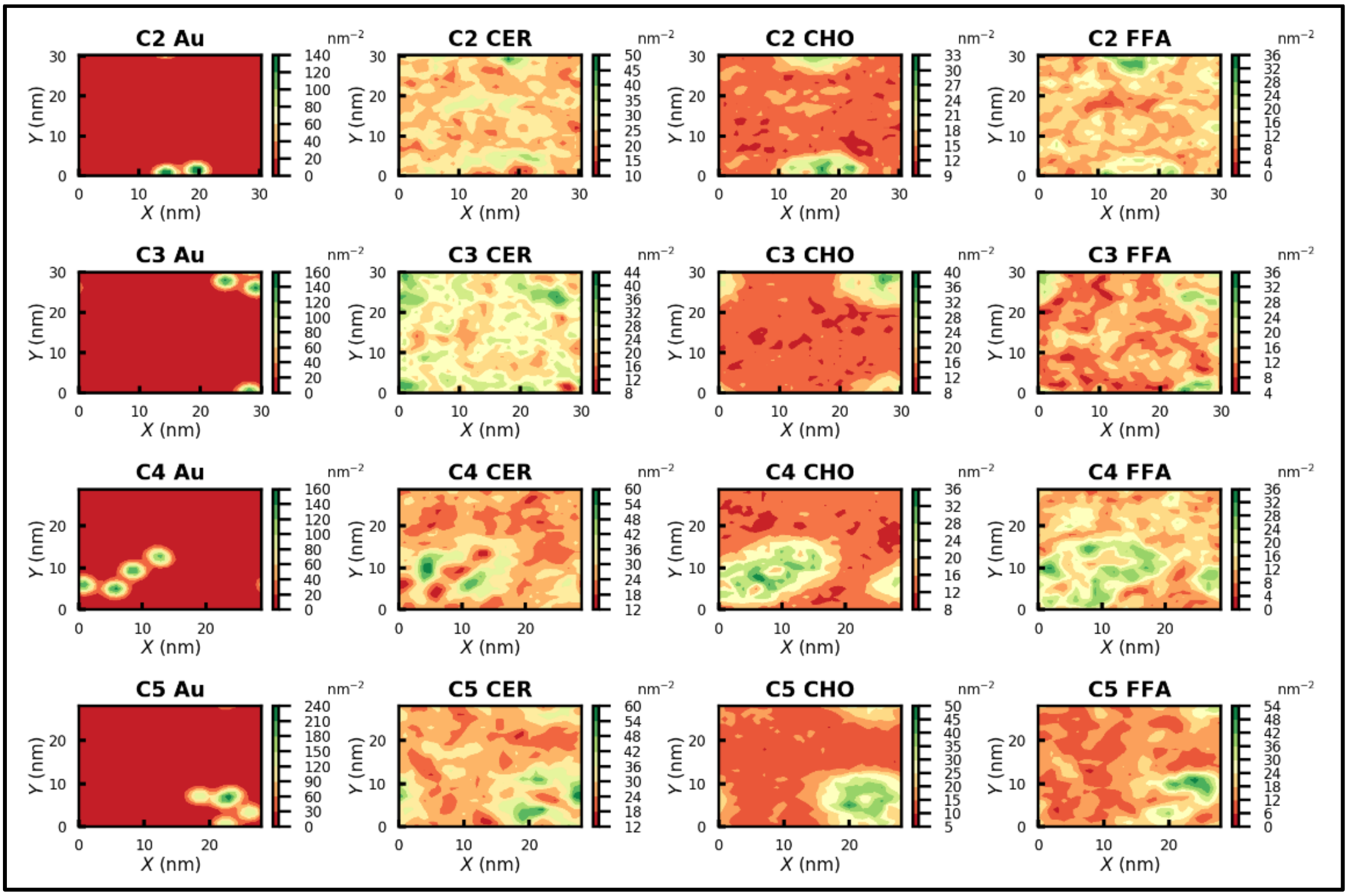


**Figure S9.** Density maps of nanoparticles, ceramide, cholesterol, and free fatty acid for C2 to C5 systems.

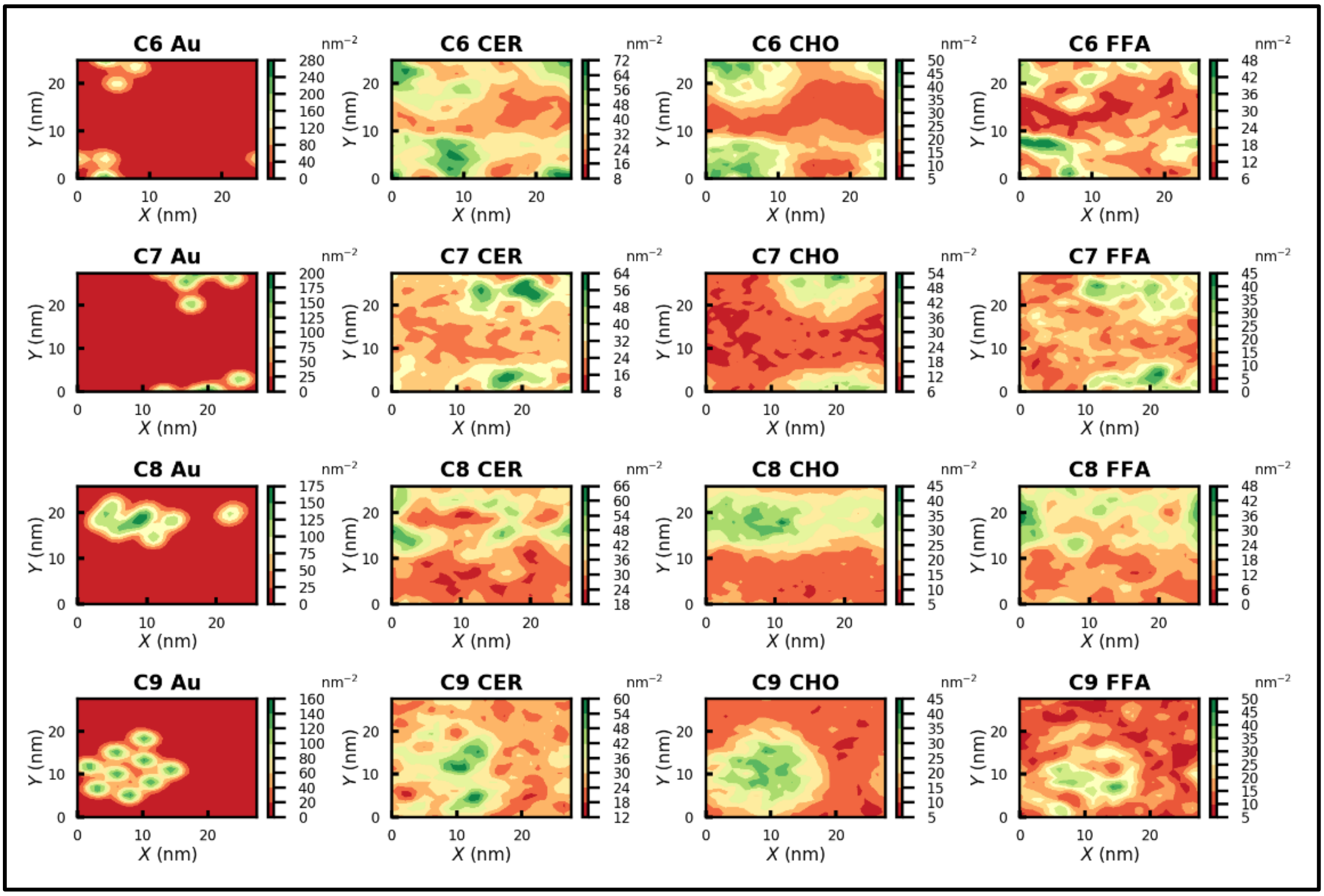


**Figure S10.** Density maps of nanoparticles, ceramide, cholesterol, and free fatty acid for C6 to C9 systems.

#### 1.8 Radial distribution function

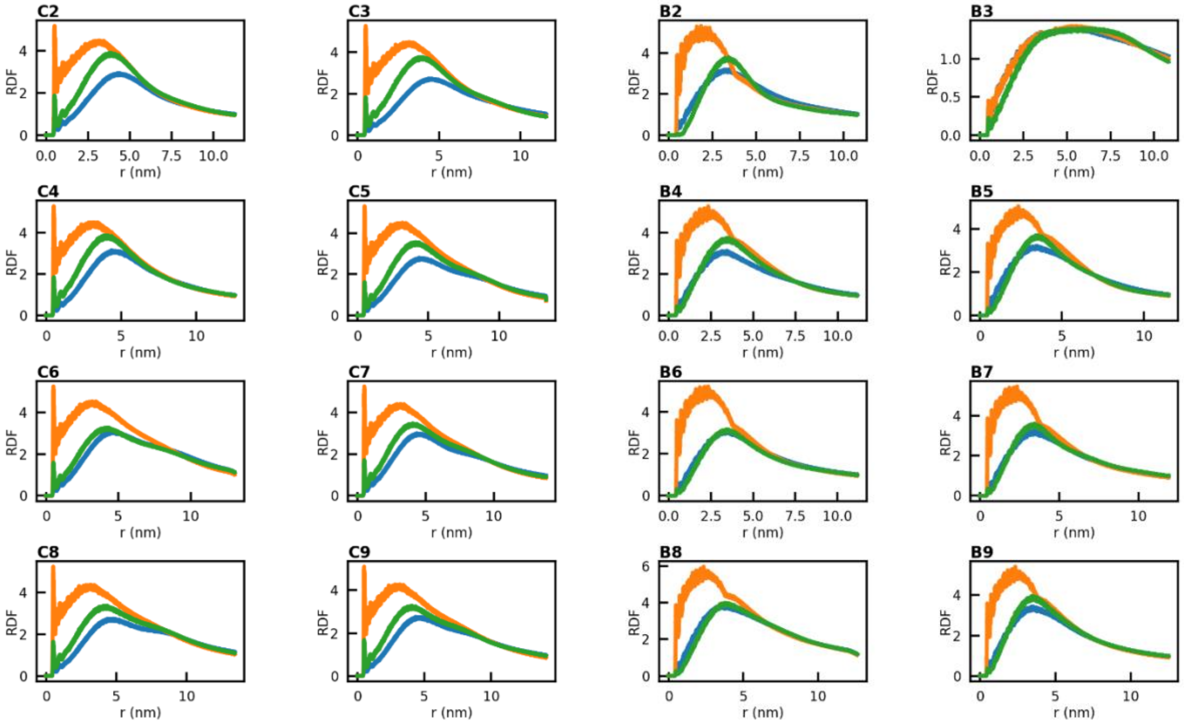


**Figure S11.** Radial distribution function of CER (green), CHO (orange) and FFA(blue) with nanoparticles.

### 2. Double membrane

#### 2.1 Snapshots

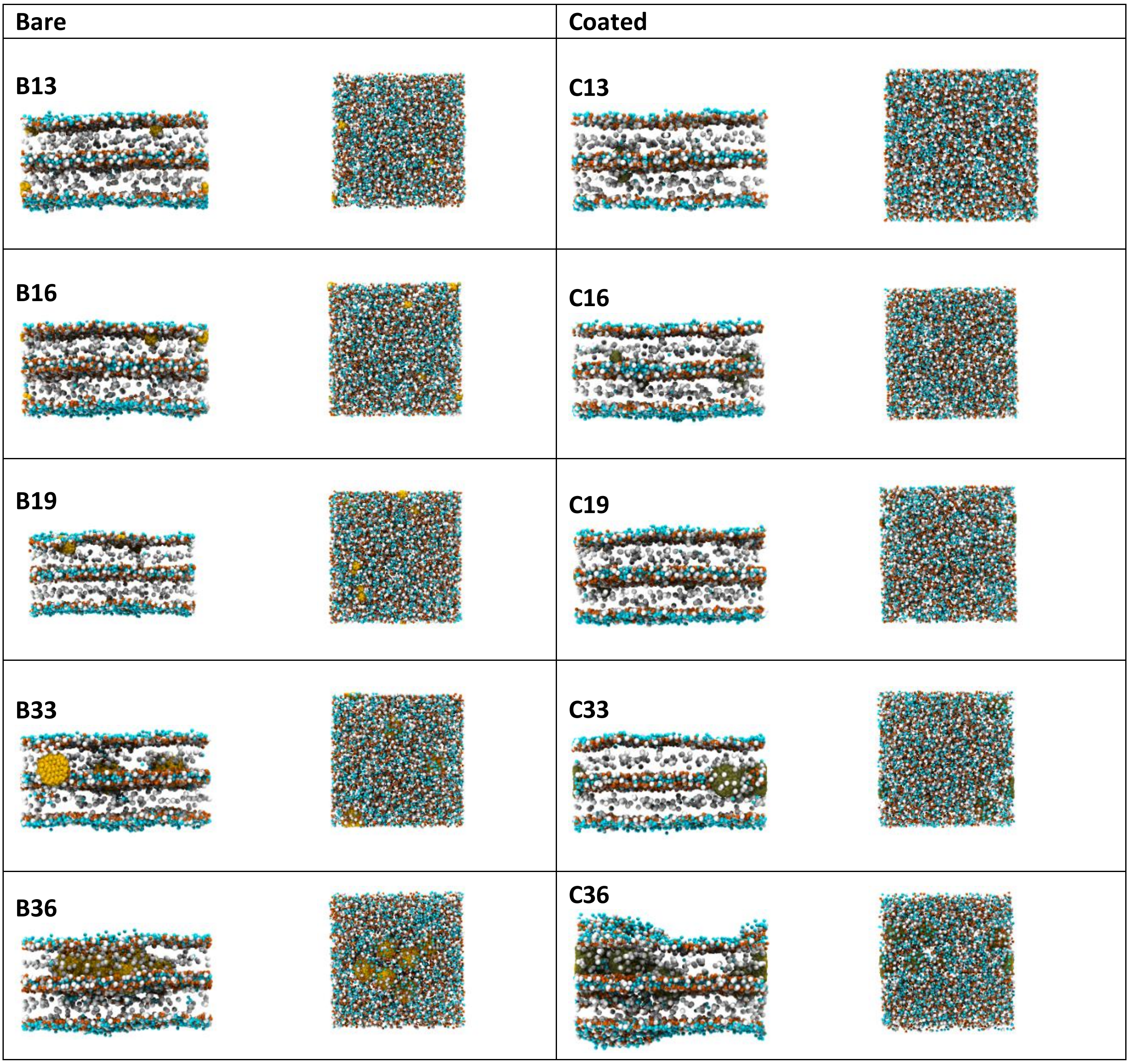


**Figure S12.** Final snapshots of double bilayer-NP system after 3 μs simulation.

#### 2.2 Trajectories

##### 2.2.1 1 μs trajectory

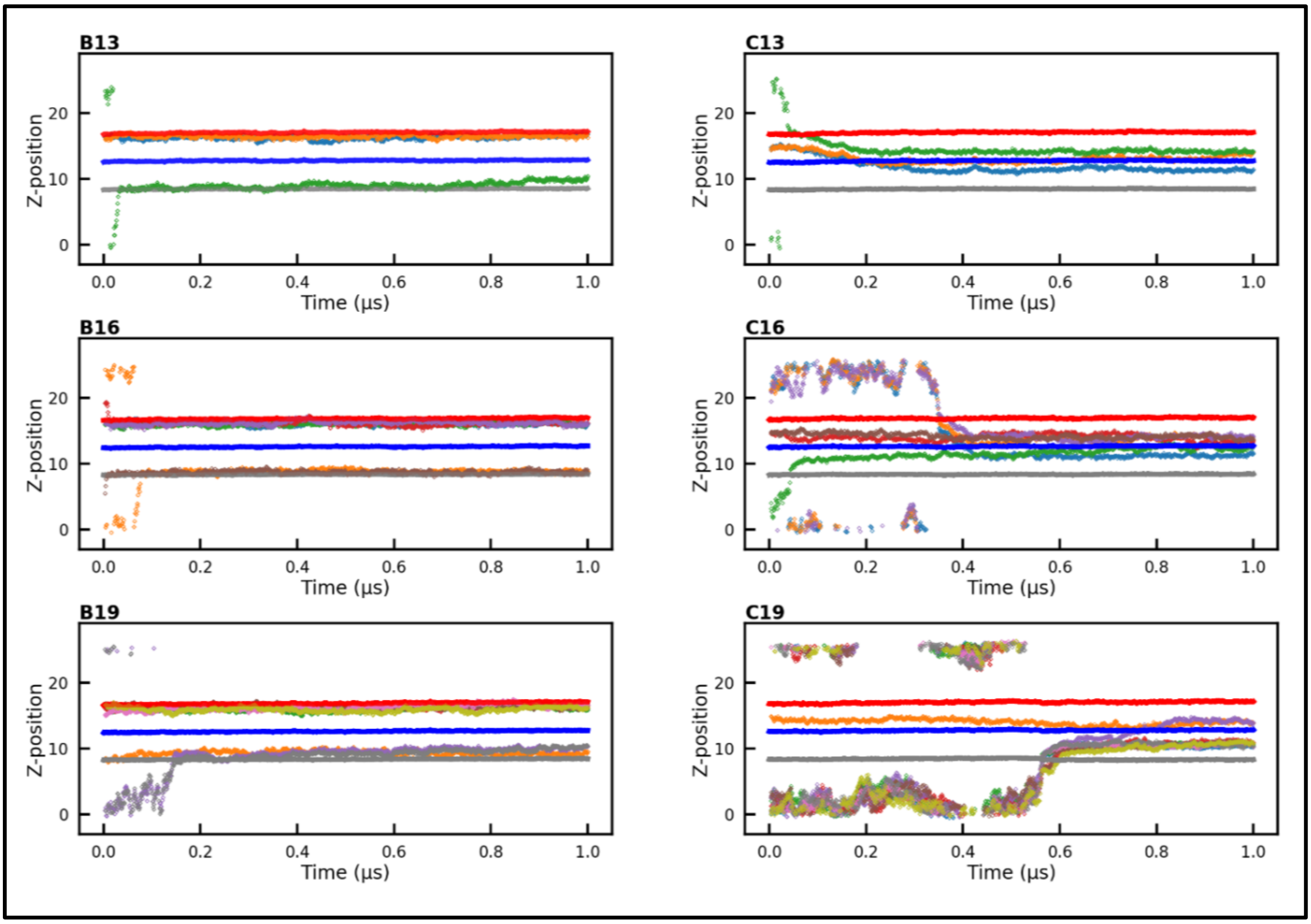


**Figure S13.** Time trajectories of double bilayer headgroup atoms and NPs (of 1 nm size) along the Z direction calculated over 1 μs simulation time. The head group data for upper, middle, and lower leaflet are shown in red, blue, and grey color respectively. Individual NP trajectories are shown in other colors.

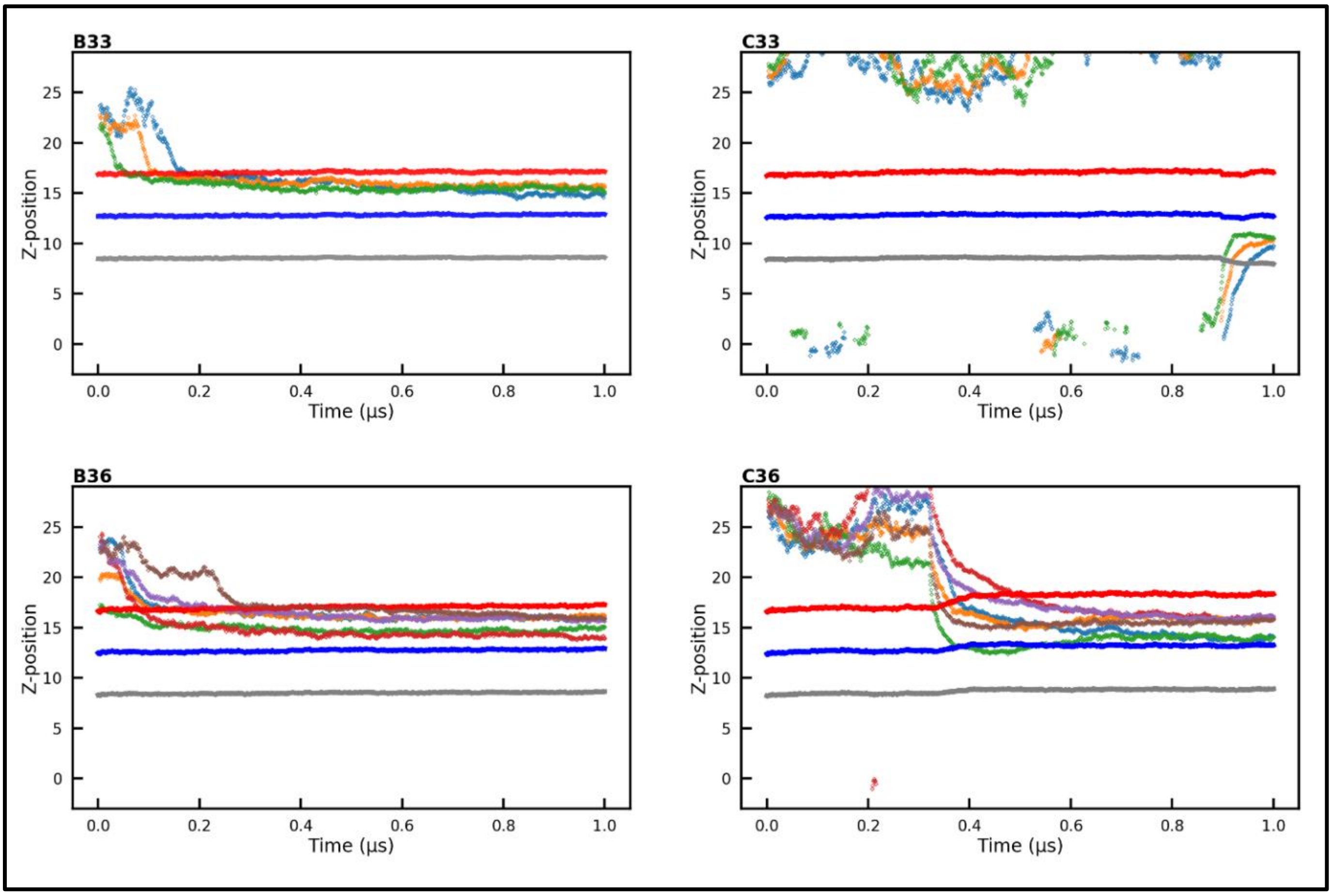


**Figure S14.** Time trajectories of double bilayer headgroup atoms and NPs (of 3 nm size) along the Z direction calculated over 1 μs simulation time. The head group data for upper, middle, and lower leaflet are shown in red, blue, and grey color respectively. Individual NP trajectories are shown in other colors.

#### 2.2.2 3 μs trajectory

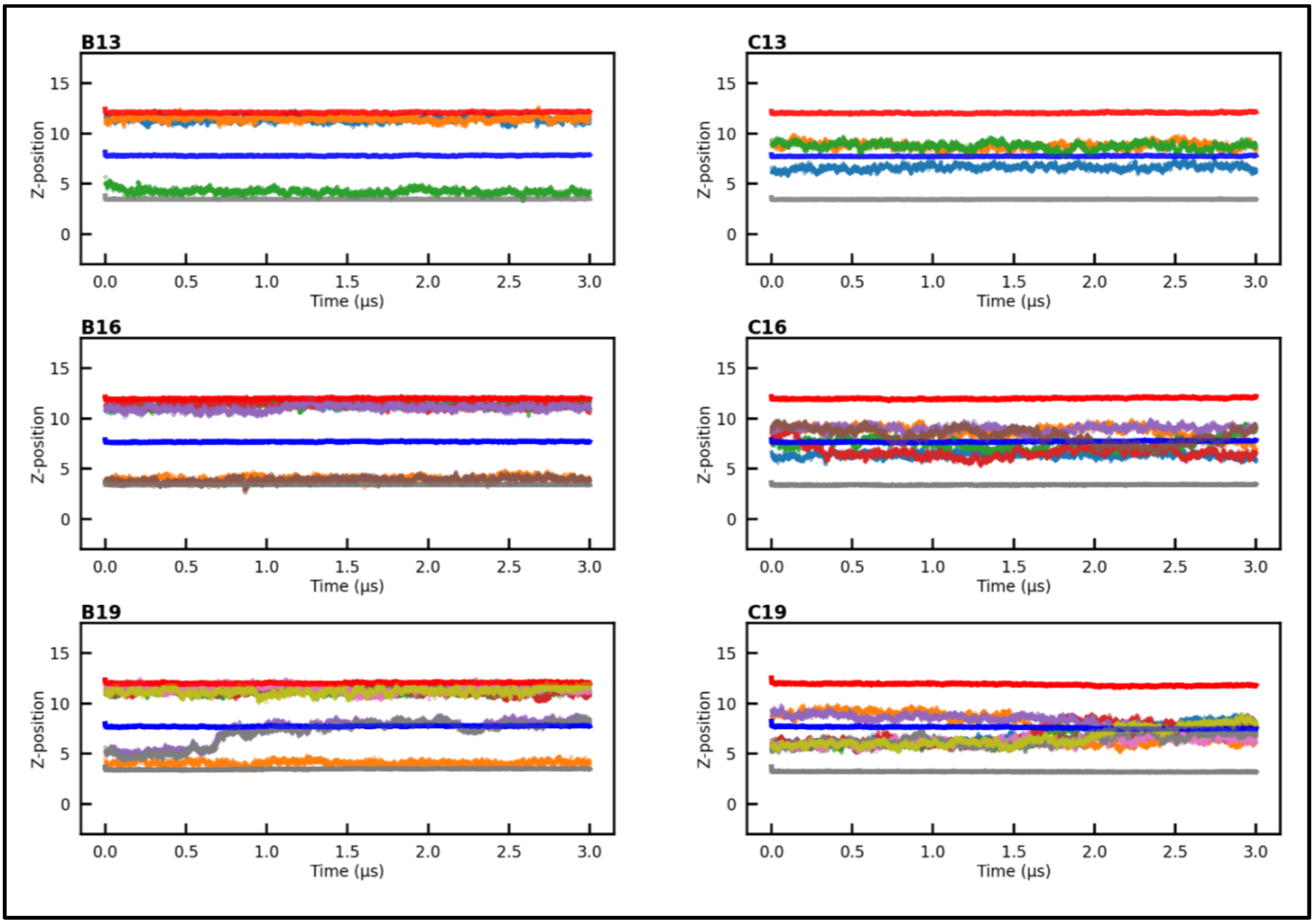


**Figure S15.** Time trajectories of double bilayer headgroup atoms and NPs (of 1 nm size) along the Z direction calculated over 3 μs simulation time. The head group data for upper, middle, and lower leaflet are shown in red, blue, and grey color respectively. Individual NP trajectories are shown in other colors.

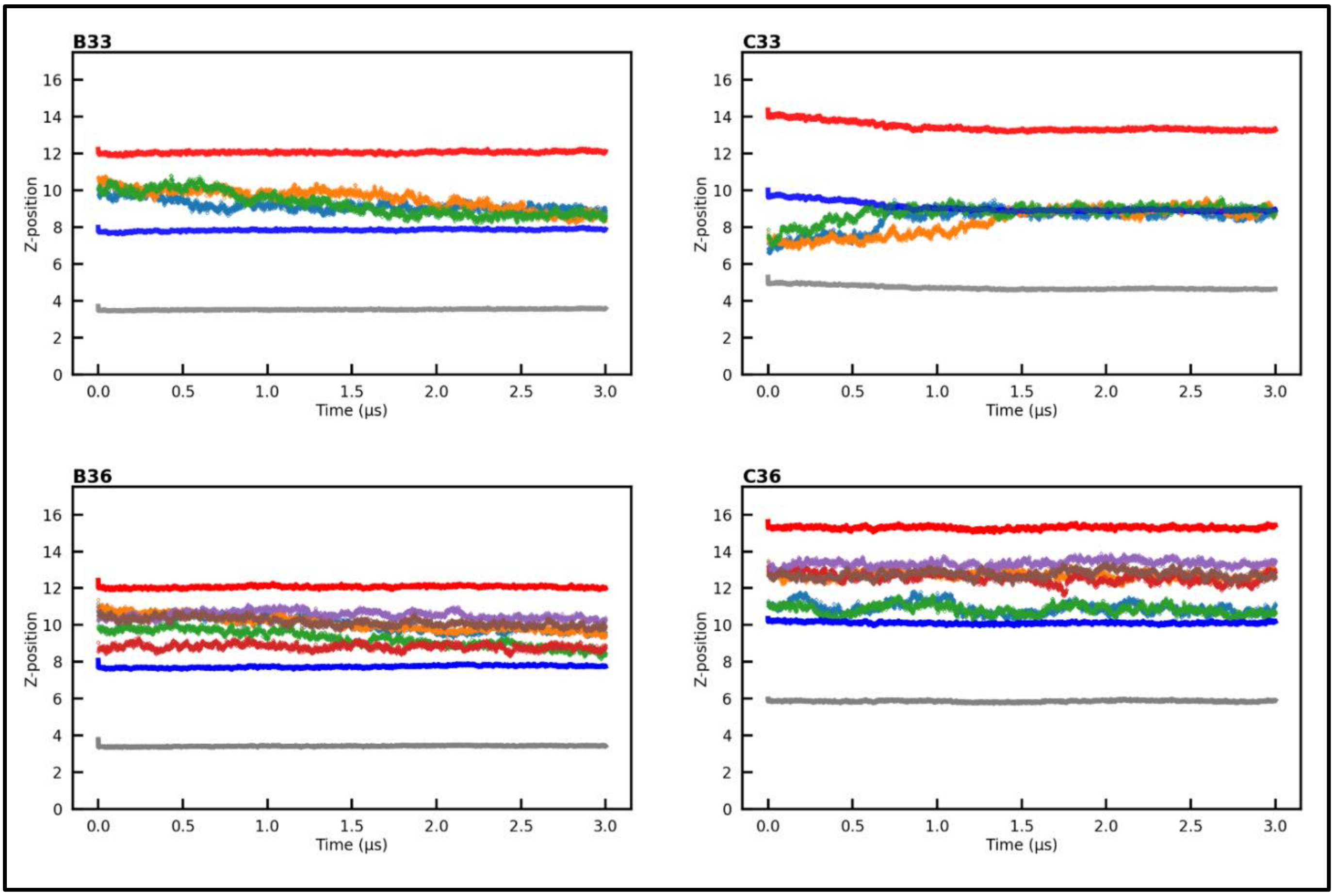


**Figure S16.** Time trajectories of double bilayer headgroup atoms and NPs (of 3 nm size) along the Z direction calculated over 1 μs simulation time. The head group data for upper, middle, and lower leaflet are shown in red, blue, and grey color respectively. Individual NP trajectories are shown in other colors.

#### 2.3 Fluctuations

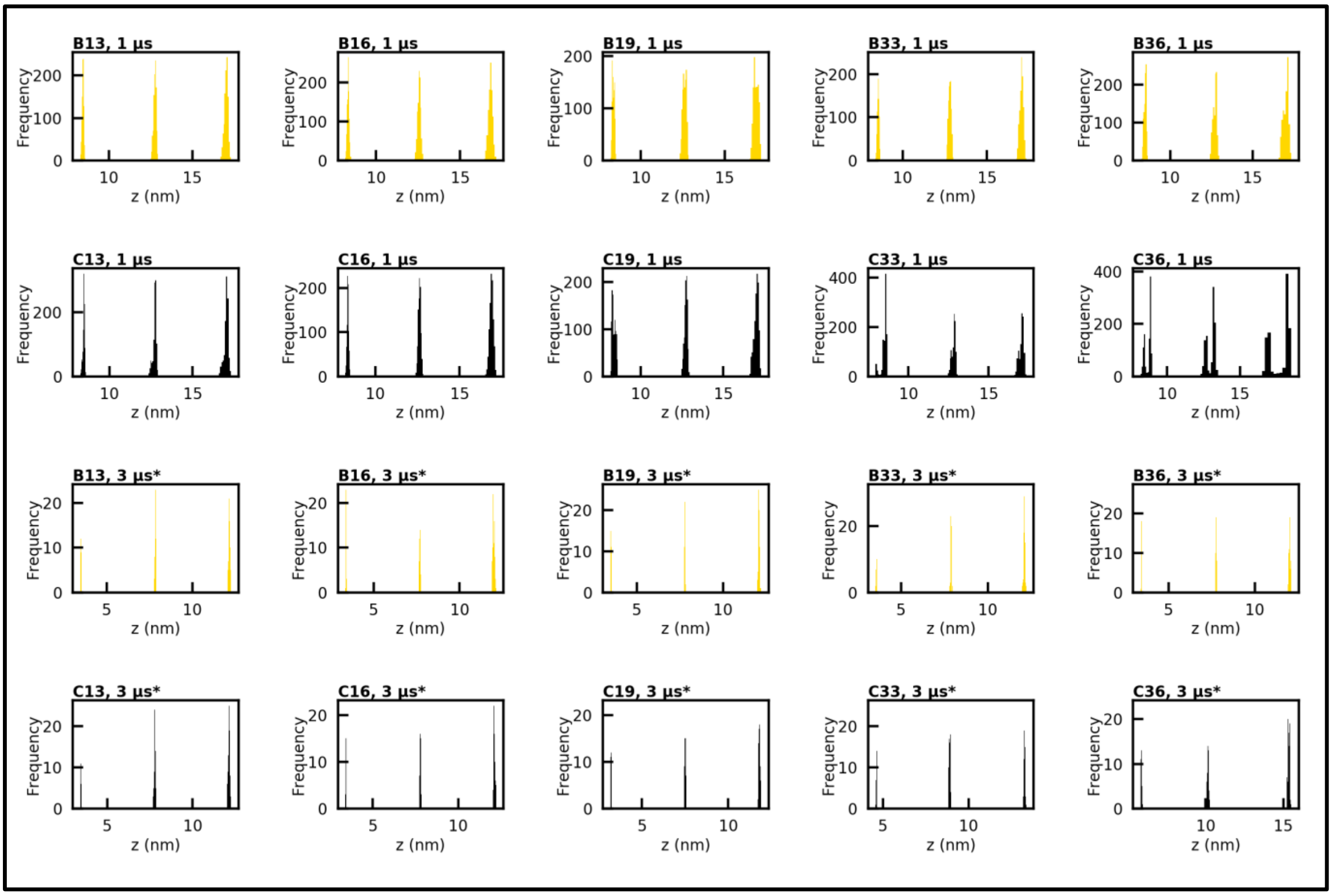

**Figure S17.** Histograms of time trajectories given in Figures S9 to S10. * Corresponds to data for last 100 ns time trajectory of 3 µs simulation. The narrower histograms for last 100 ns trajectory signify stable bilayers as compared to earlier time of simulation. Histograms for bare and coated NPs are shown in gold and black color respectively.

#### 2.4 Bilayer density along bilayer normal (z axis)

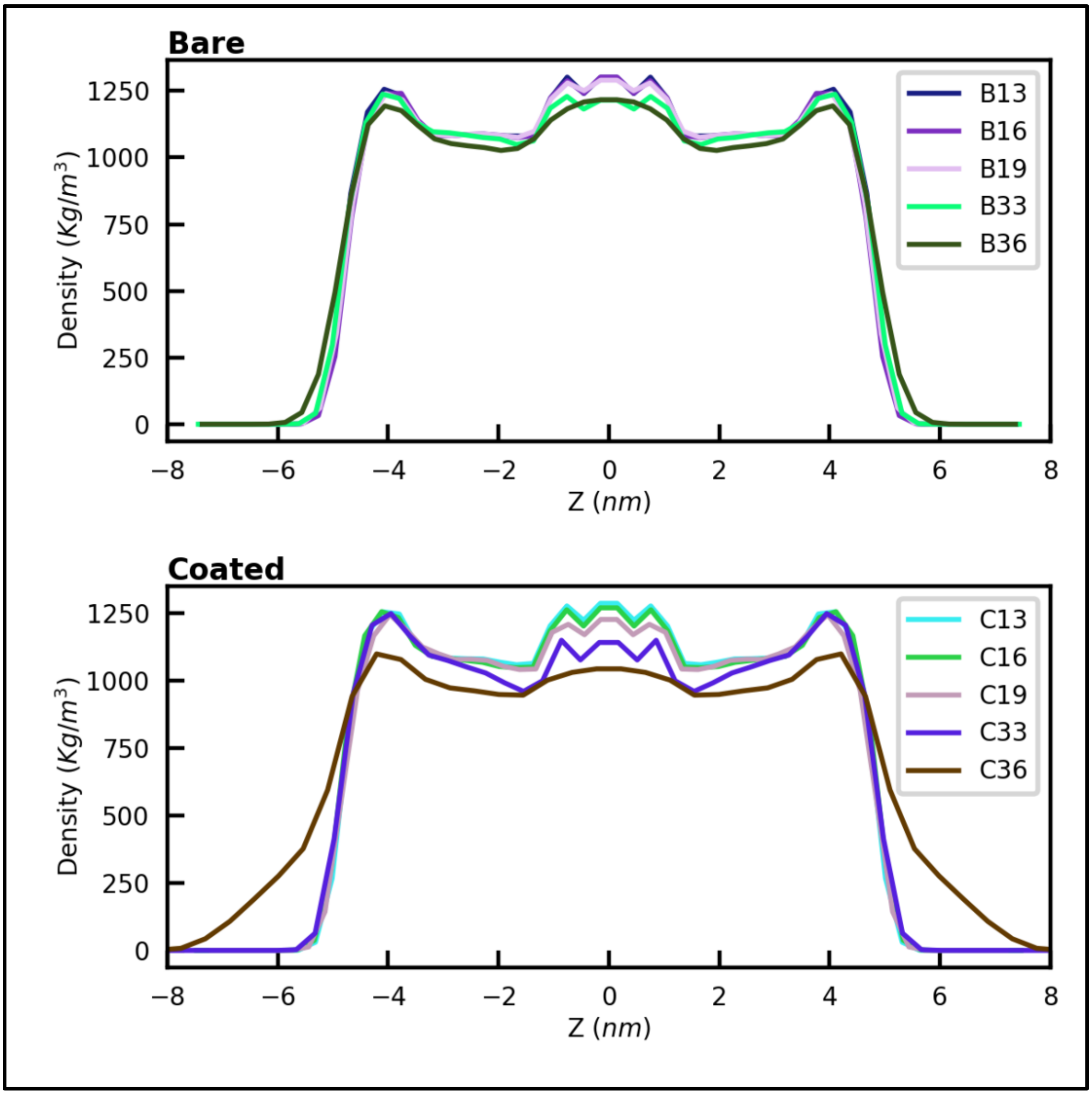


**Figure S18.** Bilayer density along z axis (last 100 ns)

#### 2.5 Lipid order parameter

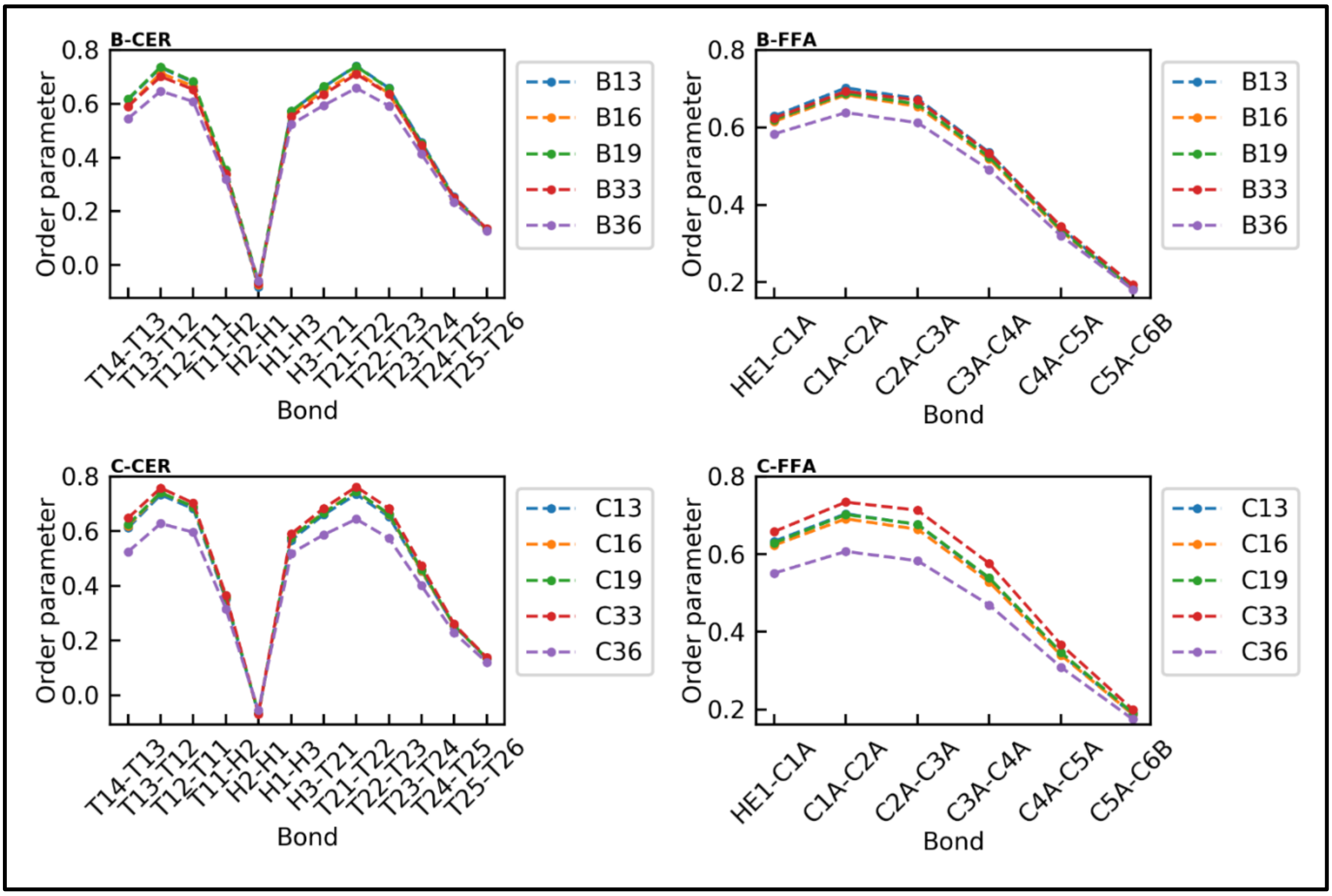


**Figure S19.** Tail order parameter of CER and FFA chains. B and C corresponds to bare and coated nanoparticle system respectively.

### 2.6 Density maps

#### 2.6.1 Bare NP, size 1 nm

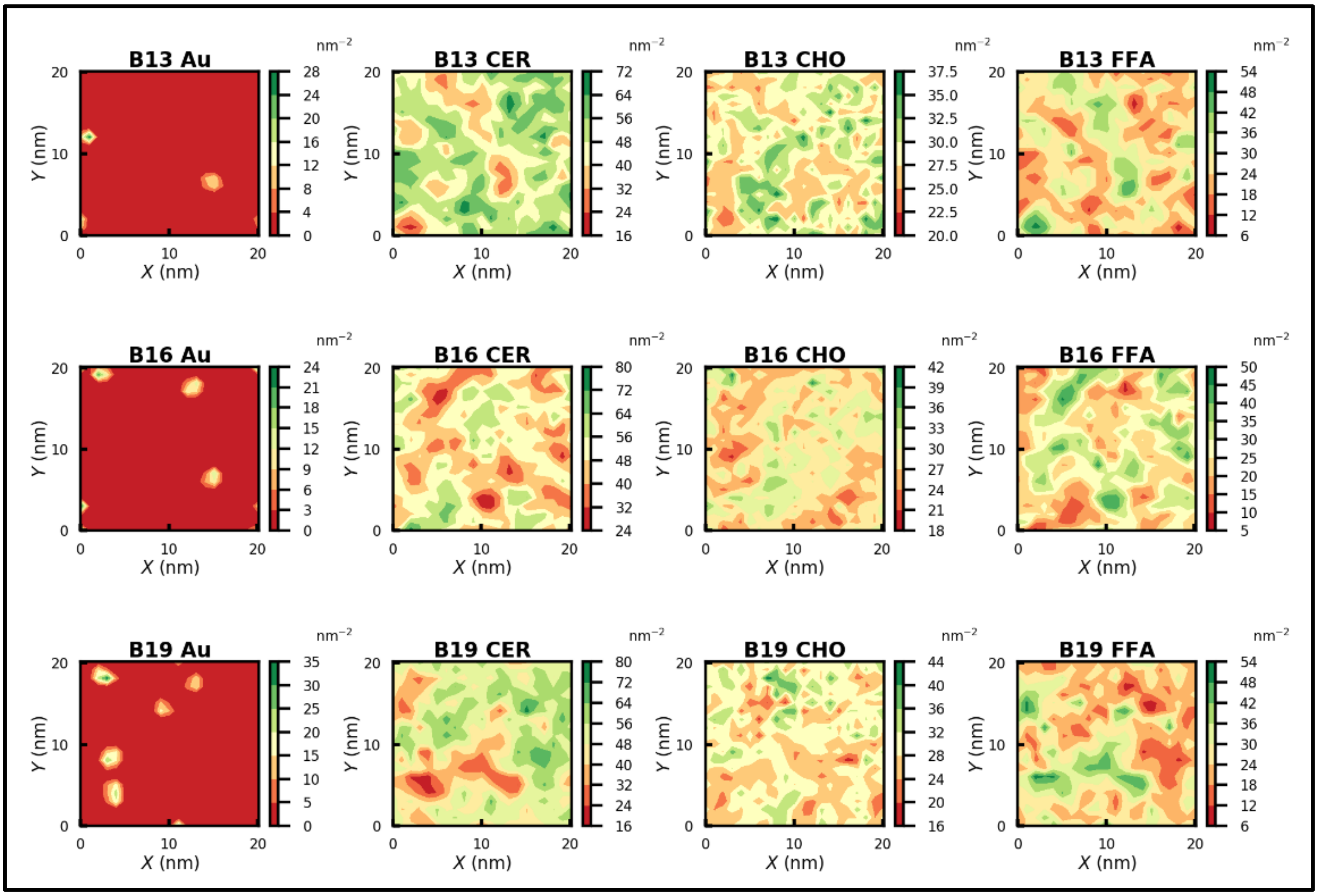


**Figure S20.** Density maps of nanoparticles, ceramide, cholesterol, and free fatty acid for B13 to B19 systems.

#### 2.6.2 Coated NP, size 1 nm

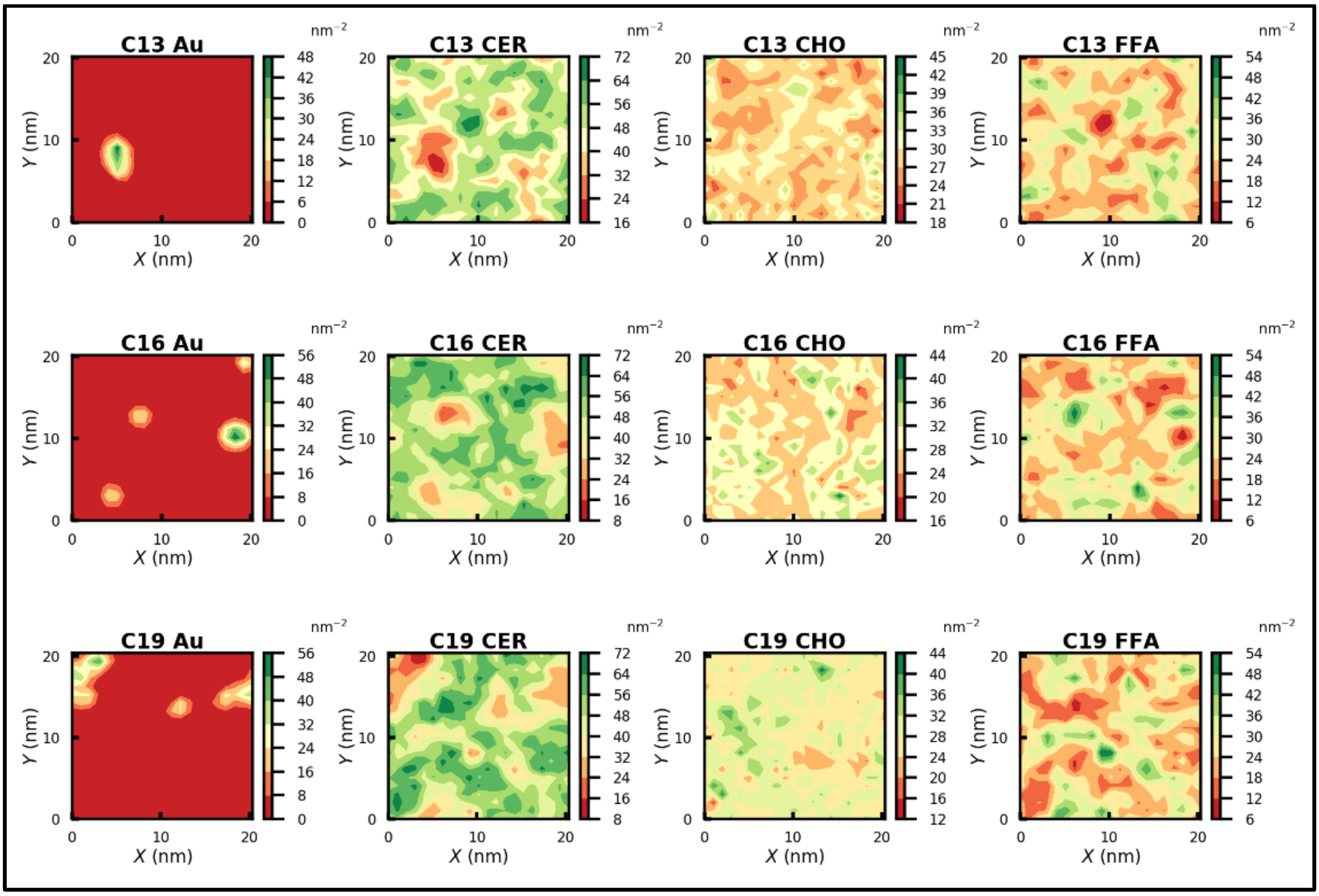


**Figure S21.** Density maps of nanoparticles, ceramide, cholesterol, and free fatty acid for C13 to C19 systems.

#### 2.6.3 Bare & Coated NP, size 3 nm

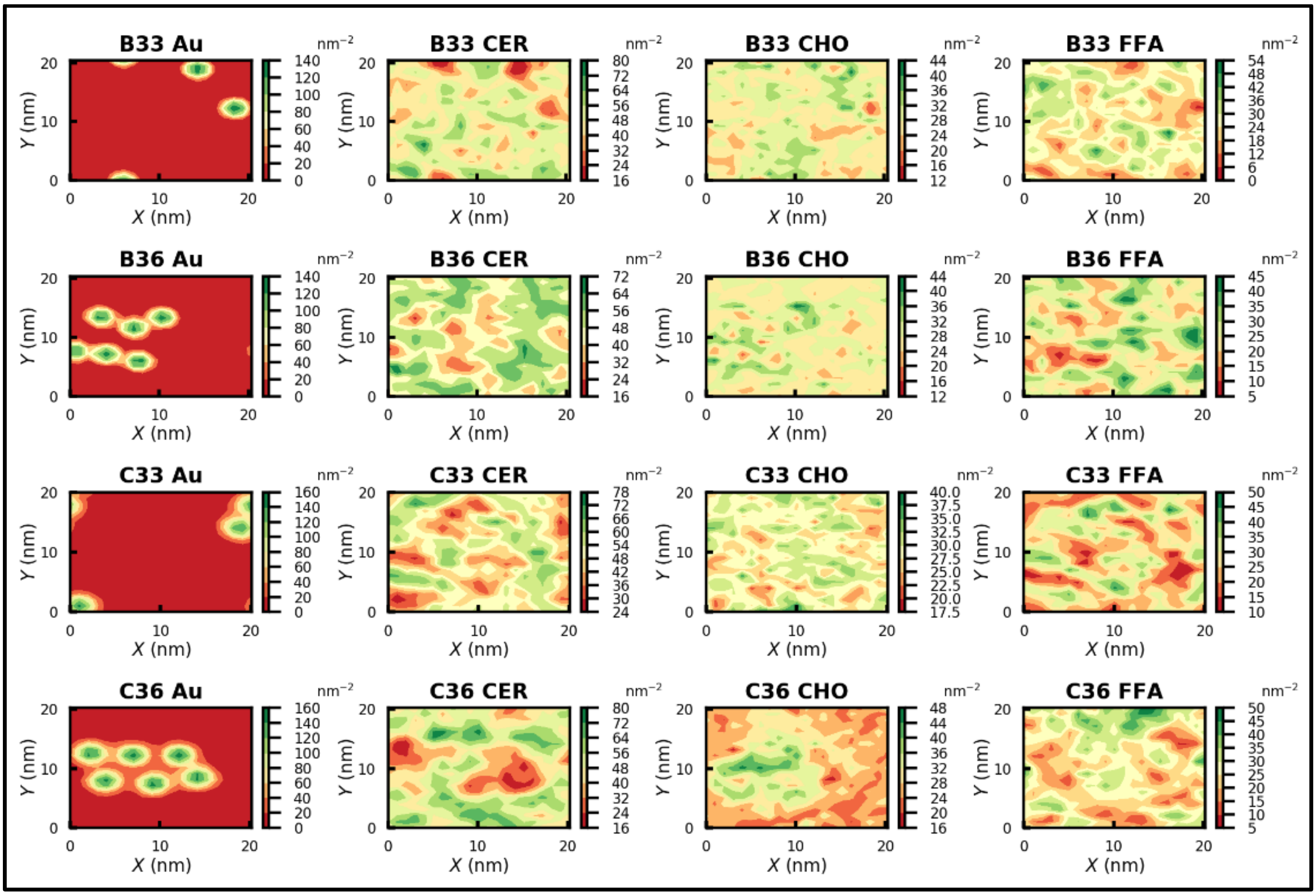


**Figure S22.** Density maps of nanoparticles, ceramide, cholesterol, and free fatty acid for B33, B36, C33 and C36 systems.

### 2.7 Radial distribuion function

#### 2.7.1 NP size 1 nm

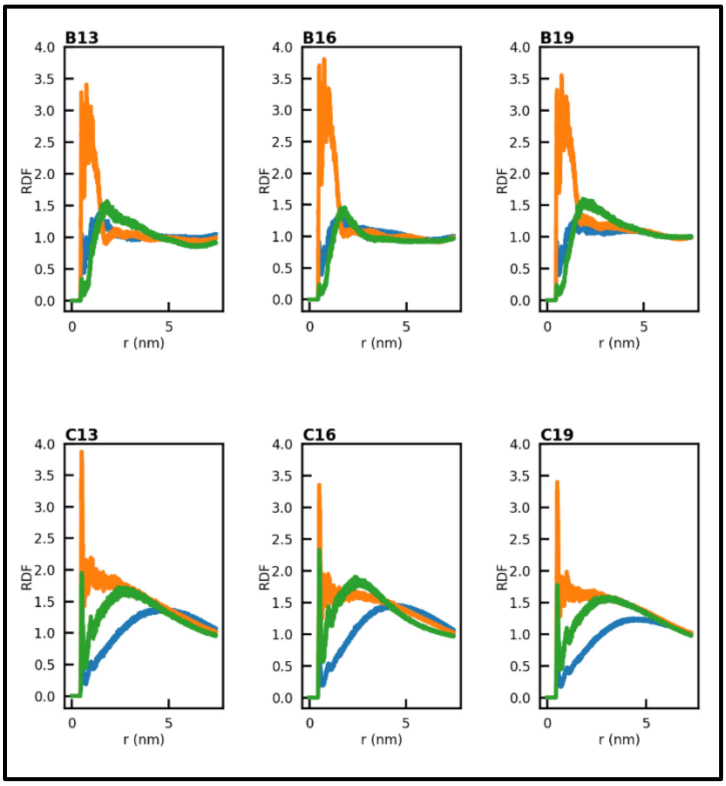


**Figure S23**. Radial distribution function of CER, CHO and FFA with nanoparticles.

#### 2.7.2 NP size 3 nm

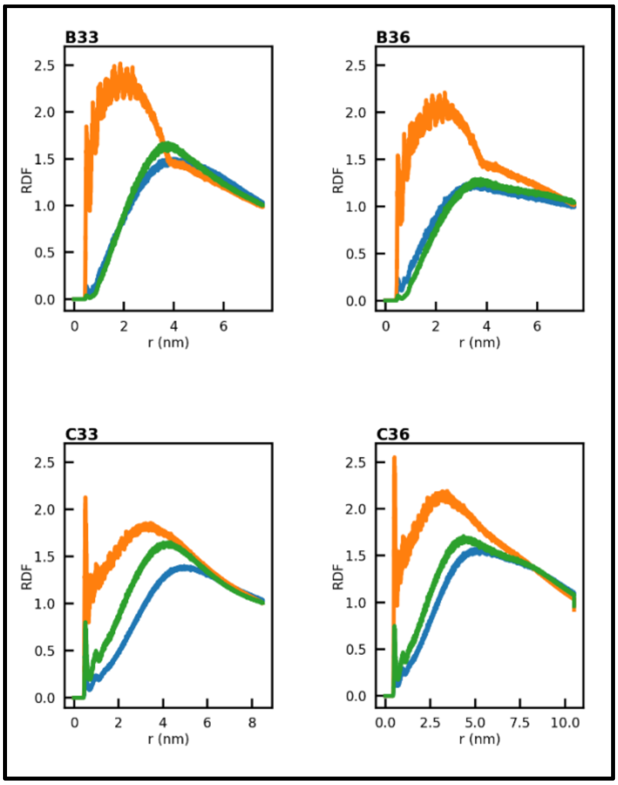


**Figure S24.** Radial distribution function of CER (green), CHO (orange) and FFA (blue) with nanoparticles
